# The MLL3/4 complexes and MiDAC act antagonistically as genome-wide regulators of H4K20ac to control a specific gene expression program

**DOI:** 10.1101/2021.12.17.473155

**Authors:** Xiaokang Wang, Wojciech Rosikiewicz, Yurii Sedkov, Baisakhi Mondal, Satish Kallappagoudar, Andrey Tvardovskiy, Richa Bajpai, Beisi Xu, Shondra M Pruett-Miller, Robert Schneider, Hans-Martin Herz

## Abstract

The mitotic deacetylase complex MiDAC has recently been shown to play a vital physiological role in embryonic development and neurite outgrowth. However, how MiDAC functionally intersects with other chromatin-modifying regulators is poorly understood. Here, we describe a physical interaction between the histone H3K27 demethylase UTX, a complex-specific subunit of the enhancer-associated MLL3/4 complexes, and MiDAC. We demonstrate that UTX bridges the association of the MLL3/4 complexes and MiDAC by interacting with ELMSAN1, a scaffolding subunit of MiDAC. Our data shows that MiDAC constitutes a negative genome-wide regulator of H4K20ac, an activity which is counteracted by the MLL3/4 complexes. MiDAC and the MLL3/4 complexes co-localize at many genomic regions, that are enriched for H4K20ac and the enhancer marks H3K4me1, H3K4me2 and H3K27ac. We find that MiDAC antagonizes the recruitment of the MLL3/4 complexes to negatively regulate H4K20ac, H3K4me2 and H3K27ac resulting in transcriptional attenuation of associated genes. In summary, our findings provide a paradigm how the opposing roles of chromatin-modifying components, such as MiDAC and the MLL3/4 complexes, balance the transcriptional output of specific gene expression programs.

## Introduction

MLL3 (also known as KMT2C) and MLL4 (also known as KMT2D) are chromatin-modifying proteins that monomethylate histone H3K4 via their catalytic SET domains. MLL3 and MLL4 exist in two separate macromolecular complexes that belong to a total of six mammalian complexes of the compositionally and functionally highly conserved COMPASS family (Cenik & Shilatifard 2020, Herz 2016, Shilatifard 2012). All complexes share identical core subunits but also contain complex-specific subunits that are conserved only within one of three metazoan branches. Each metazoan branch is represented by two mammalian complexes, namely the SET1A/B complexes (branch one), the MLL1/2 complexes (branch two) and the MLL3/4 complexes (branch three). UTX (also known as KDM6A) exists as a complex-specific subunit within the MLL3/4 complexes and acts as a histone H3K27 demethylase removing methyl groups from the inhibitory histone marks H3K27me3 and H3K27me2 via its Jumonji C domain (Hong et al 2007, Lee et al 2013). Our previous studies and the work of others have demonstrated that the MLL3/4 complexes function as major H3K4 monomethyltransferases on enhancers, providing a model in which prior removal of H3K27me3/2 via UTX is required on inactive or poised enhancers, before they can transition to an activated state via addition of H3K4me1 through MLL3/4 (Herz et al 2010, Herz et al 2012, Hu et al 2013, Lee et al 2013, Rickels et al 2017). Despite our increasing understanding of UTX, MLL3 and MLL4 in regulating enhancer activity in development and disease, many functional aspects of the MLL3/4 complexes remain ill-defined.

Lysine residues on histones can also be acetylated by histone acetyltransferases (HATs) and deacetylated by histone deacetylases (HDACs) (Shahbazian and Grunstein, 2007). Histone acetylation has been found on all four core histones (H1-4) and can be deposited by specific writers and recognized by site-specific readers at multiple lysine residues on each core histone (Barnes et al., 2019; Gräff and Tsai, 2013; Marmorstein and Zhou, 2014). The mammalian genome encodes multiple classes of histone deacetylases, among which the class I histone deacetylases (HDAC1- 3, 8) are the most well studied (Milazzo et al., 2020). HDAC1-3 are assembled into large multi-subunit protein complexes to regulate the acetylation state of histones and other non-histone proteins. The integration of HDAC1-3 into these scaffolds both strongly enhances the enzymatic activities and also determines the specificity of these HDAC complexes (Banks et al., 2020; Bantscheff et al., 2011; Millard et al., 2013, 2016; Turnbull et al., 2020; Wang et al., 2020; Watson et al., 2012). HDAC1/2 are integrated into the NuRD, SIN3, CoREST and MiDAC complexes, while HDAC3 is a component of the SMRT/NCoR complex (Laherty, 1997; Li, 2000; Oberoi, 2011; Xue, 1998) (reviewed in (Millard et al 2017).

The mitotic deacetylase complex MiDAC, which is comprised of the scaffolding subunits DNTTIP1, ELMSAN1 (also known as MIDEAS) and the histone deacetylases HDAC1 or HDAC2, was initially identified in a chemoproteomic screen as being specifically enriched on a HDAC-inhibitor-bound resin in cells stalled in G2/prophase of mitosis following nocodazole treatment (Bantscheff et al., 2011). TRERF1 and ZNF541 are paralogs of ELMSAN1 with a more tissue-specific expression pattern and have also been described as scaffolding subunits of MiDAC (Bantscheff et al 2011, Choi et al 2008, Hao et al 2011). MiDAC exhibits a tetrameric architecture with each monomer consisting of DNTTIP1, ELMSAN1 and HDAC1 or HDAC2 thus resembling a three-dimensional X-shape with the HDAC1/2 catalytic sites at the four extremities of the complex suggesting that MiDAC may simultaneously target multiple nucleosomes and may be highly processive (Itoh et al., 2015; Turnbull et al., 2020). MiDAC components are located predominantly in the soluble nuclear fraction throughout the cell cycle and loss of MiDAC function causes misalignment of chromosomes in metaphase (Turnbull et al., 2020). Furthermore, mutations of DNTTIP1 and ELMSAN1 have been identified in different cancer types (Cheng, 2017; Sawai, 2018; SW Piraino, 2017; Xu, 2018; Zhang, 2018). MiDAC also constitutes an important regulator of a neural gene expression program to ensure proper neuronal maturation and/or neurite outgrowth during neurogenesis (Mondal et al., 2020). Homozygous knock-out mouse embryos lacking either DNTTIP1 or ELMSAN1 die at day E16.5 with severe anemia and a clear malformation of the heart (Turnbull et al., 2020).

Despite recent advances in our understanding of MiDAC’s structure and physiological role, it is unknown how MiDAC integrates into chromatin regulatory complexes and pathways. Applying a large scale interactome analysis we describe here we also describe an association of UTX with MiDAC. We show that UTX and ELMSAN1 form the interface between MiDAC and the MLL3/4 complexes. We demonstrate that the loss of MiDAC function results in a genome-wide increase of H4K20ac, suggesting that H4K20ac constitutes a MiDAC substrate *in vivo*. The genome-wide increase of H4K20ac in *Dnttip1* KO mouse embryonic stem cells (mESCs) coincides with increased UTX and MLL4 occupancy on many genomic elements, indicating that MiDAC negatively affects the recruitment of the MLL3/4 complexes to chromatin. While H3K4me1 enrichment is slightly decreased, H3K4me2 enrichment is higher at regions displaying increased UTX and MLL4 occupancy, suggesting an activity switch of MLL3/4 towards H3K4me2 in the presence of H4K20ac or absence of MiDAC function. However, in *Mll3/4* double knockout (DKO) mESCs we observe an increase of H3K27me3 at regions with decreased H4K20ac and lower DNTTIP1 occupancy, indicating that H3K27me3 may be involved in suppressing H4K20ac, despite reduced MiDAC recruitment. Taken together, our study reveals for the first time a functional intersection between MiDAC and the MLL3/4 complexes in regulating H4K20ac and describes the antagonistic relationship of these chromatin-modifying complexes in their role to properly balance transcription of a specific gene expression program.

## Results

### The mitotic deacetylase complex MiDAC associates with the MLL3/4 complexes

To identify novel candidates that regulate the function of the MLL3/4 complexes, we immunoprecipitated UTX, a H3K27 demethylase and complex-specific subunit of the MLL3/4 complexes, from nuclear extracts of human embryonic kidney cells (HEK293 cells). Mass spectrometry (MS) analysis detected as expected all components of the MLL3/4 complexes including all core subunits, complex-specific subunits and the H3K4 methyltransferases MLL3 and MLL4 (Figure 1A; Table S1). Additionally, we also identified two novel and strong UTX interactors: DNTTIP1 and ELMSAN1 (Figure 1A; Table S1). The association of UTX with DNTTIP1 and ELMSAN1 along with other subunits of the MLL3/4 complexes was confirmed by Western blotting (WB) (Figure 1B). Reciprocal immunoprecipitations (IPs) of ELMSAN1 and DNTTIP1 from nuclear extracts of HEK293 cells confirmed an interaction of these proteins with components of the MLL3/4 complexes including UTX, MLL3/4 and the core subunits RBBP5 and ASH2L (Figure 1C; Figure S1). Consistent with previous studies showing that the histone deacetylases HDAC1 and HDAC2 interact with DNTTIP1 and ELMSAN1 to be integrated into the histone deacetylase complex MiDAC (Bantscheff et al 2011, Mondal et al 2020, Turnbull et al 2020), both our ELMSAN1 and DNTTIP1 IPs also purified HDAC1 and HDAC2 (Figure 1C; Figure S1). An association of HDAC1 and HDAC2 with UTX was also observed following UTX IP providing further evidence that UTX associates indeed with MiDAC (Figure1B; Table S1). Glycerol gradient fractionation of FLAG-affinity purified UTX from nuclear extracts of HEK293 cells confirmed that MiDAC co-migrates with the MLL3/4 complexes (Figure 1D, red box). A substantial amount of MiDAC also associated with UTX outside the MLL3/4 complexes indicating that not all overexpressed UTX is incorporated into the MLL3/4 complexes and that MiDAC might directly interact with UTX (Figure 1D, blue box). The high number of ELMSAN1 peptides in our UTX IP, as assessed by mass spectrometry analysis and the fact that after UTX IP a significant amount of MiDAC co-migrated with UTX outside the MLL3/4 complexes, suggests that UTX might constitute a bridging component between MiDAC and the MLL3/4 complexes (Figure 1D; Table S1). Both ELMSAN1 and its paralog TRERF1, but not the paralog ZNF541, are expressed in HEK293 cells (data not shown). Thus, due to a potential redundancy between ELMSAN1 and TRERF1 we generated an *ELMSAN1 TRERF1* double knockout (DKO) HEK293 cell line to establish the interaction interface between MiDAC and MLL3/4 complexes in more detail (Figure 1E; Figure S2). In *ELMSAN1 TRERF1* DKO cells, DNTTIP1 protein is almost completely lost (Figure 1E). Interestingly, reexpression of either FLAG-ELMSAN1 or FLAG-TRERF1 in *ELMSAN1 TRERF1* DKO cells resulted in stabilization of DNTTIP1, suggesting that MiDAC scaffolding subunits need to be integrated into MiDAC to retain their stability (Figure 1E). Reexpression of FLAG-DNTTIP1, FLAG-ELMSAN1, or FLAG-TRERF1 in *ELMSAN1 TRERF1* DKO cells followed by FLAG IPs, showed that only IPs from FLAG-ELMSAN1 expressing cells were able to purify UTX, while FLAG-DNTTIP1 and FLAG-TRERF1 expressing cells could not (Figure 1E). To test whether UTX was able to interact with MiDAC outside of the MLL3/4 complexes, we utilized HCT116 colorectal carcinoma cells which contain a homozygous frameshift mutation in *MLL3* (*MLL3-/- MLL4+/+*) (Watanabe et al 2011). We employed CRISPR/Cas9 genome editing to also obtain HCT116 cells that are wild-type (WT) for *MLL3* (*MLL3+/+ MLL4+/+*) or contain a promoter deletion within the *MLL4* locus (*MLL3-/- MLL4-/-*) (Figure S3A and B). Transient transfection of FLAG-UTX into these HCT116 cell lines followed by FLAG IPs showed that UTX interaction with the MiDAC members DNTTIP1, ELMSAN1 and HDAC1 did not depend on MLL3/4 or core subunits of the MLL3/4 complexes such as RBBP5 and WDR5 suggesting a direct interaction between UTX and MiDAC (Figure S3C). In conclusion, our data imply that UTX functions as a bridging factor within the MLL3/4 complexes to mediate their association with MiDAC via its scaffolding subunit ELMSAN1 (Figure 1F).

**Figure 1.**
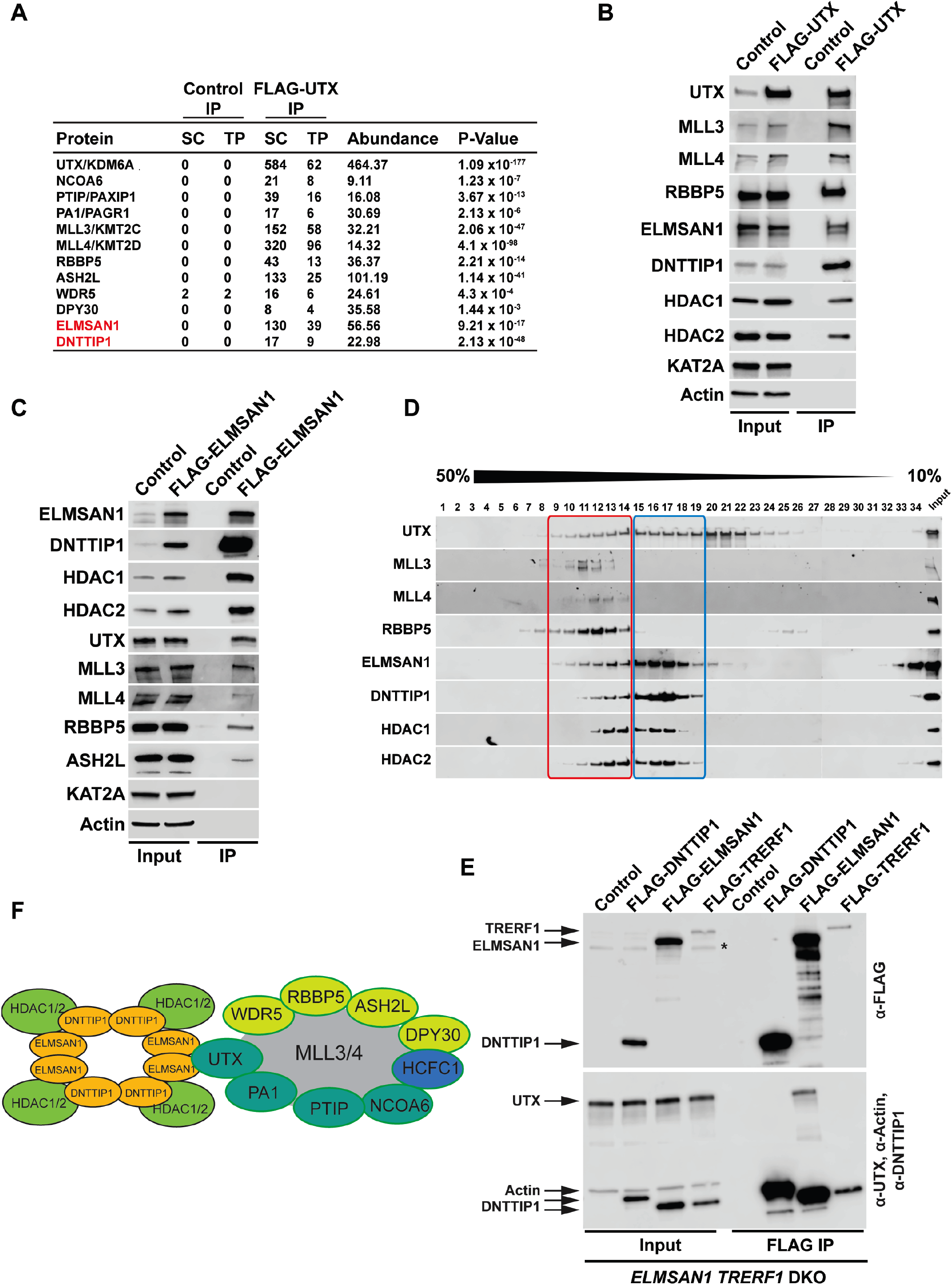
The MLL3/4 complexes associate with the mitotic deacetylase complex (MiDAC). **(A)** FLAG-UTX immunoprecipitation (IP) followed by mass spectrometry (MS) identifies all known subunits of the MLL3/4 complexes along with ELMSAN1, and DNTTIP1. SC – spectral counts; TP – peptide counts; abundance = SC x 50 (kDa)/protein size (kDa). **(B)** Western blot (WB) of FLAG-UTX IP from HEK293 cells confirming interaction of UTX with ELMSAN1, DNTTIP1, HDAC1, and HDAC2. UTX interacts with the H3K4 methyltransferases MLL3 and MLL4, RBBP5 a core component of the MLL3/4 complexes, ELMSAN1, DNTTIP1, HDAC1, and HDAC2, but does not interact with KAT2A a histone acetyltransferase. HEK293 cells with a FLAG-tag expressing plasmid were used as an IP control. Nuclear extracts were used as input. Actin was used as a loading control for the inputs. **(C)** WB of FLAG-ELMSAN1 IP from HEK293 cells confirming interaction of ELMSAN1 with the MiDAC components DNTTIP1, HDAC1, HDAC2 and members of the MLL3/4 complexes including UTX, the H3K4 methyltransferases MLL3 and MLL4, and the two core components RBBP5 and ASH2L. KAT2A was used as a negative control. HEK293 cells with a FLAG-tag expressing plasmid were used as an IP control. Nuclear extracts were used as input. Actin was used as a loading control for the inputs. **(D)** Glycerol gradient sedimentation after FLAG-UTX IP reveals co-fractionation of the MiDAC subunits ELMSAN1, DNTTIP1, HDAC1, and HDAC2 with components of the MLL3/4 complexes (red box). **(E)** WB of FLAG IP from *ELMSAN1 TRERF1* double knockout (*ELMSAN1 TRERF1* DKO) HEK293 cells expressing FLAG-DNTTIP1, FLAG-ELMSAN1, or FLAG-TRERF1. Among the MiDAC subunits, only ELMSAN1 interacts with UTX. Reexpression of FLAG-ELMSAN1 or FLAG-TRERF1 in *ELMSAN1 TRERF1* DKO cells results in restoration of DNTTIP1 levels. * marks a nonspecific band. **(F)** Diagram showing that ELMSAN1 and UTX form the nexus between MiDAC and the MLL3/4 complexes.

### MiDAC is a genome-wide negative regulator of H4K20ac

We have previously shown by WB that a loss of MiDAC function in mESCs results in a bulk increase of H4K20ac (Mondal et al., 2020). To confirm these findings by an antibody-independent more quantitative method, we applied LC-MS/MS and found significantly higher levels of H4K20ac in two independent *Dnttip1* KO clones compared to WT mESCs (Figure S4). To investigate site specific changes in H4K20ac profiles genome-wide, we used ChIP-seq to profile H4K20ac in WT and *Dnttip1* KO mESCs. Additionally, ChIP-seq was also carried out for H3K4me1/2/3, H3K27ac, and H3K27me3, UTX, and MLL4 in the same cell lines. Reanalysis of our previously published DNTTIP1 ChIP-seq data set from WT and *Dnttip1* KO mESCs (Mondal et al., 2020) identified a total of 38,572 DNTTIP1 binding sites (FC<0.5, FDR<0.05) (Figure 2A). Consistent with the bulk increase of H4K20ac in *Dnttip1* KO mESCs (Figure S4) (Mondal et al., 2020), we observe higher levels of H4K20ac on the majority of DNTTIP1-bound genomic regions in *Dnttip1* KO compared to WT mESCs (Figure 2A and B). We also detected a milder but higher enrichment of H3K4me2 and H3K4me3 without considerable alterations of H3K4me1 on regions that are targeted by MiDAC (DNTTIP1-bound regions) in *Dnttip1* KO versus WT mESCs (Figure 2C). Figure 2D depicts two enhancer regions and the promoter of the *Spry4* locus as an individual example (Figure 2D, red boxes). All three regulatory elements are bound by MiDAC (DNTTIP1) in WT mESCs and display a strong increase of H4K20ac upon *Dnttip1* KO, which is accompanied by higher enrichment of H3K4me2/me3 (Figure 2D, red boxes). Additionally, we also observe a concomitant increase of H3K27ac and lower enrichment of the repressive H3K27me3 mark while H3K4me1 is not significantly changed (Figure 2D, red boxes). In summary, our findings indicate that MiDAC constitutes a genome-wide negative regulator of H4K20ac by inhibiting the accumulation of H4K20ac on promoters and putative enhancers.

**Figure 2.**
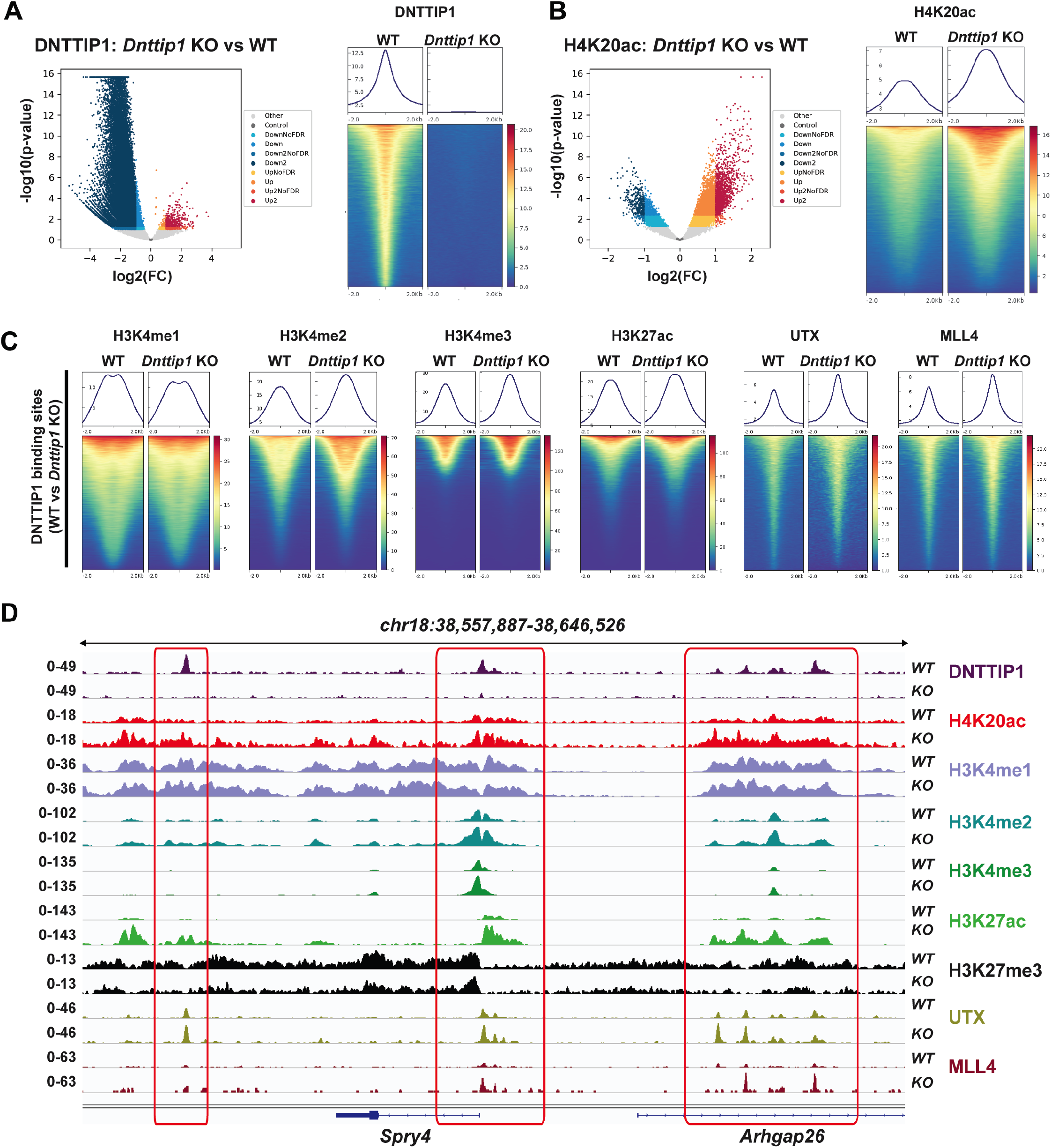
MiDAC is a genome-wide negative regulator of H4K20ac. **(A-B)** Volcano plots (left) and heatmaps (right) showing the genome-wide loss of DNTTIP1 (A) and genome-wide increase of H4K20ac (B) in *Dnttip1* KO versus WT mouse embryonic stem cells (mESCs). Heatmaps in (A) and (B) are centered on 38,572 identified DNTTIP1 peaks by comparing DNTTIP1 occupancy between WT and *Dnttip1* KO mESCs (FC>2). **(C)** Heatmaps centered on the 38,572 DNTTIP1 peaks identified in WT compared to *Dnttip1* KO mESCs in (A). Occupancy of H3K4me1, H3K4me2, H3K4me3, H3K27ac, UTX, and MLL4 in WT and *Dnttip1* KO mESCs is displayed. **(D)** Genome browser tracks depicting the ChIP-seq profiles of DNTTIP1, H4K20ac, H3K4me1, H3K4me2, H3K4me3, H3K27ac, H3K27me3, UTX, and MLL4 in WT and *Dnttip1* KO mESCs at the promoter and two enhancer regions of the *Spry4* locus.

### MiDAC antagonizes the function of the MLL3/4 complexes on chromatin

To further dissect the functional relationship between MiDAC and the MLL3/4 complexes we wanted to know how the loss of MiDAC function would affect chromatin occupancy of members of the MLL3/4 complexes. We found that a much higher fraction of UTX-bound or MLL4-bound regions displayed increased rather than decreased binding of UTX or MLL4 in *Dnttip1* KO compared to WT mESCs (Figure 3A). In total, we identified 4,826 peaks with higher UTX and 26,084 peaks with higher MLL4 occupancy in *Dnttip1* KO mESCs (both FC>2) (Figure 3A). Interestingly, the DNTTIP1 binding sites identified in Figure 2C also displayed increased UTX and MLL4 occupancy in *Dnttip1* KO mESCs. Occupancy of UTX and MLL4 on putative enhancers and the promoter of the *Spry4* locus was also elevated in *Dnttip1* KO compared to WT mESCs (Figure 2C and D, red boxes). Additionally, regions with upregulated UTX binding (FC>2) in *Dnttip1* KO mESCs also showed an increase in MLL4, H4K20ac, H3K4me2, and H3K27ac enrichment (Figure S5A). Similarly, regions with upregulated MLL4 binding (FC>2) in *Dnttip1* KO mESCs displayed increased UTX, H4K20ac, H3K4me2, and H3K27ac enrichment (Figure S5B). To assess the relationship between MiDAC and the MLL3/4 complexes more stringently, we subsequently only focused on the 1,283 regions that displayed both increased UTX and MLL4 occupancy (FC>2) in *Dnttip1* KO mESCs (Figure S5C). These UTX/MLL4 upregulated regions also showed increased enrichment of H4K20ac, H3K4me2, and H3K27ac in *Dnttip1* KO versus WT mESCs (Figure 3B). Although the MLL3/4 complexes have been shown to catalyze both H3K4me1 and H3K4me2 on enhancers (Hu et al 2013, Lee et al 2013), we did not observe a significant change of H3K4me1 enrichment across the regions that showed both increased UTX and MLL4 occupancy in *Dnttip1* KO mESCs (Figure 3B). Similarly, no difference in H3K4me1 enrichment was detected on DNTTIP1-bound regions or individual promoter or putative enhancer elements of the *Spry4* locus in *Dnttip1* KO compared to WT mESCs (Figure 2C and D). Instead, in all cases H3K4me2 was increased (Figure 2C and D, and 3B). Based on this observation we also identified the regions with increased H3K4me2 enrichment (FC>2) in *Dnttip1* KO mESCs (2,396 regions) and observed higher enrichment of UTX, MLL4, H4K20ac, and H3K27ac without major changes in H3K4me1 on these sites in *Dnttip1* KO mESCs (Figure 3C). Interestingly, both promoter or putative enhancer regions with either increased H4K20ac or H3K4me2 enrichment or UTX/MLL4 occupancy tend to be associated with upregulated gene expression in *Dnttip1* KO compared to WT mESCs (Figure 3D). Taken together, this suggests that MiDAC antagonizes the recruitment and/or activity of the MLL3/4 complexes at promoters and putative enhancers by regulating H4K20ac and H3K4me2 without significantly affecting H3K4me1.

**Figure 3.**
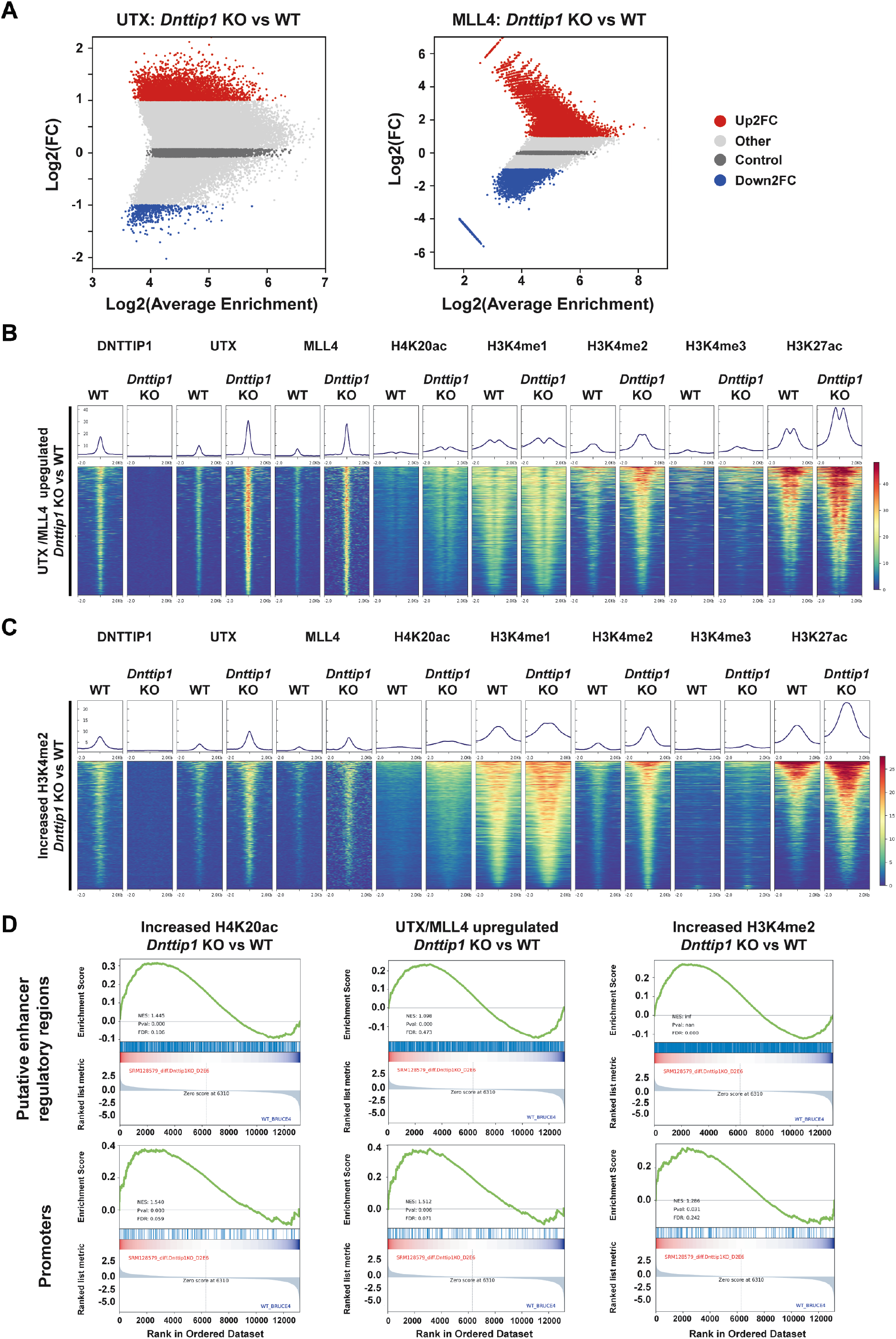
MiDAC negatively regulates the occupancy of the MLL3/4 complexes at regulatory elements genome-wide. **(A)** MA plots displaying the log_2_ fold change of UTX (left) and MLL4 (right) occupancy in *Dnttip1* KO compared to WT mESCs. A significantly higher number of UTX and MLL4 upregulated peaks (FC>2) than downregulated peaks (FC>2) is detected in *Dnttip1* KO versus WT mESCs. **(B)** Heatmaps centered on 1,283 UTX/MLL4 upregulated peaks (both FC>2) in *Dnttip1* KO compared to WT mESCs. Occupancy of DNTTIP1, UTX, MLL4, H4K20ac, H3K4me1, H3K4me2, H3K4me3, and H3K27ac in WT and *Dnttip1* KO mESCs is displayed. **(C)** Heatmaps centered on 2,396 regions with upregulated H3K4me2 (FC>2) in *Dnttip1* KO versus WT mESCs. Occupancy of DNTTIP1, UTX, MLL4, H4K20ac, H3K4me1, H3K4me2, H3K4me3, and H3K27ac in WT and *Dnttip1* KO mESCs is displayed. **(D)** Gene set enrichment analysis (GSEA) showing transcriptional upregulation of genes associated with regions of increased H4K20ac (left), UTX/MLL4 upregulated peaks (middle), and regions of increased H3K4me2 (right) in *Dnttip1* KO versus WT mESCs.

### MiDAC and the MLL3/4 complexes oppose each other’s function to balance a jointly regulated gene expression program

To better understand whether the relationship between MiDAC and the MLL3/4 complexes is based on reciprocity we performed ChIP-seq to profile DNTTIP1, H4K20ac, H3K4me1, H3K4me2, H3K4me3, H3K27ac, and H3K27me3 in WT and *Mll3/4* double knockout (DKO) mESCs (Figure S6) (Dorighi et al., 2017). Unexpectedly, most DNTTIP1-bound sites displayed decreased DNTTIP1 enrichment in *Mll3/4* DKO mESCs (Figure 4A). Altogether, we identified 10,462 sites with reduced DNTTIP1 occupancy (FC<0.5) in *Mll3/4* DKO compared to WT mESCs (Figure 4A). Interestingly however, we observed that a majority of regions with detectable H4K20ac showed a significant decrease of H4K20ac enrichment (5,513 sites) (FC<0.5) in *Mll3/4* DKO mESCs (Figure 4B). Likewise, this reduction in H4K20ac along with a reduction in H3K4me1, H3K4me2, H3K4me3, and H3K27ac was also detected at the 10,462 regions with reduced DNTTIP1 occupancy in *Mll3/4* DKO mESCs (Figure 4C). H3K27me3 inversely correlated with the aforementioned histone marks showing an increase both at sites with decreased DNTTIP1 occupancy and regions with lower H4K20ac enrichment in *Mll3/4* DKO mESCs (Figure 4C). Thus, the detected changes in histone modification patterns between *Mll3/4* DKO and *Dnttip1* KO mESCs particularly as it pertains to H4K20ac, H3K4me2, H3K27ac, and H3K27me3 anticorrelate at regions that are co-regulated by MiDAC and the MLL3/4 complexes (Figure 2C and D, 3B and C, 4C). This general finding is also obvious at putative enhancers regions and the promoter of the *Spry4* gene locus where we found H4K20ac, H3K4me2, H3K4me3, and H3K27ac to be increased in *Dnttip1* KO mESCs but reduced in *Mll3/4* DKO mESCs while H3K27me3 was reduced in *Dnttip1* KO mESCs but elevated in *Mll3/4* DKO mESCs (Figure 4D). This anticorrelative behavior as exemplified on the *Spry4* locus at the level of specific histone marks (Figure 4D) also aligned well with the gene expression changes we observed in *Dnttip1* KO compared to *Mll3/4* DKO mESCs. For example, promoters or putative enhancers with decreased enrichment of H4K20ac in *Mll3/4* DKO mESCs were strongly associated with transcriptional repression (Figure S7) while promoters and putative enhancers with increased H4K20ac in *Dnttip1* KO mESCs displayed a significantly enhanced tendency for increased gene expression (Figure 3D). Furthermore, *Spry4* transcription was increased in *Dnttip1* KO but reduced in *Mll3/4* DKO mESCs (Figure 5A and Figure S8). Transcriptome-wide we found a similar trend. Genes that were upregulated in the absence of MiDAC function showed decreased expression in *Mll3/4* DKO mESCs while downregulated genes in *Dnttip1* KO mESCs displayed increased expression in Mll3/4 DKO mESCs (Figure 5B). Overall, this suggests that the MLL3/4 complexes and MiDAC act antagonistically as genome-wide regulators of H4K20ac to transcriptionally balance a jointly regulated gene expression program.

**Figure 4.**
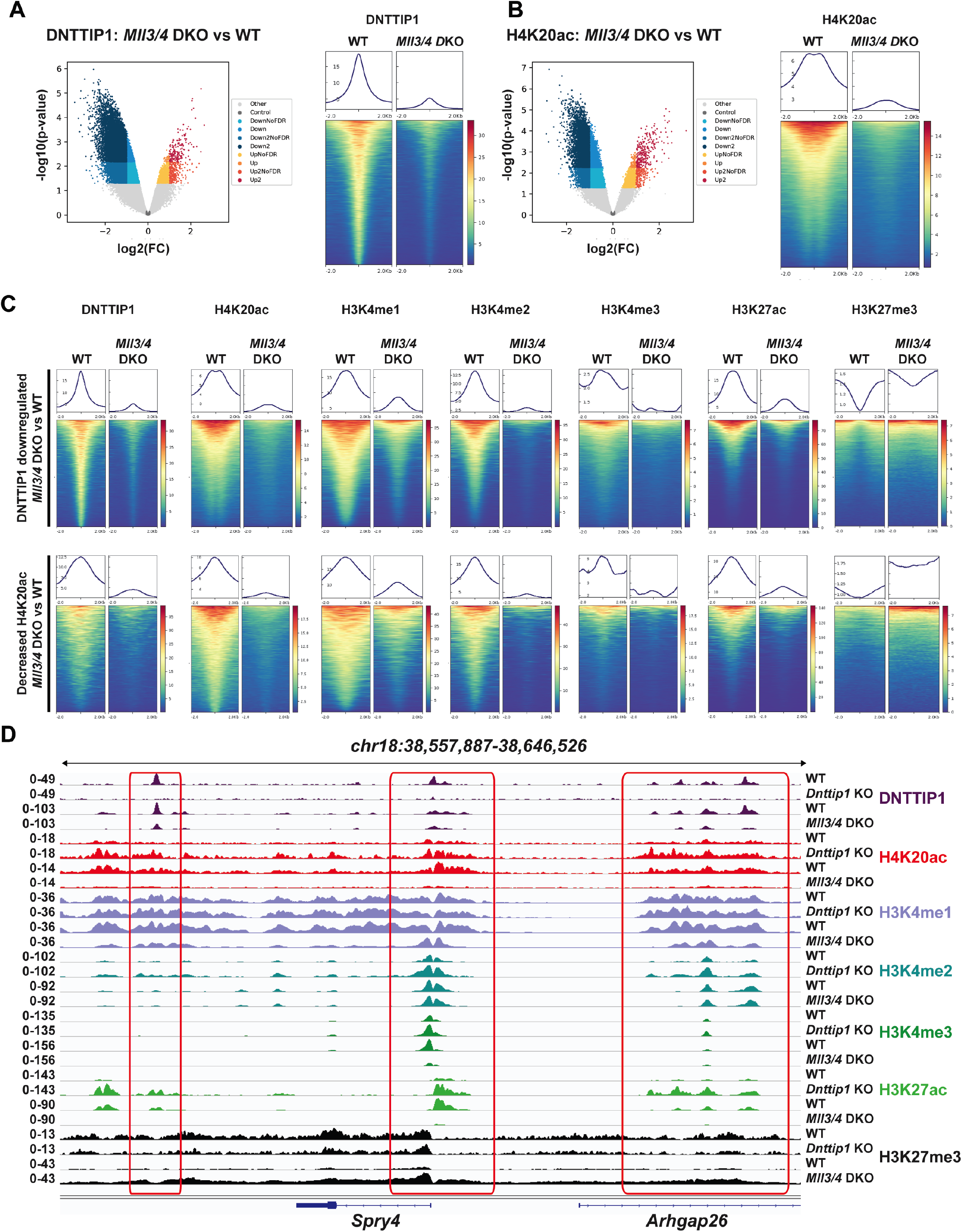
MiDAC and the MLL3/4 complexes act antagonistically as genome-wide regulators of H4K20ac. **(A-B)** Volcano plots (left) and heatmaps (right) showing the genome-wide reduction of DNTTIP1 (A) and genome-wide decrease of H4K20ac (B) in *Mll3/4* DKO compared to WT mESCs. Heatmaps in (A) and (B) are centered on 10,462 DNTTIP1 peaks with a FC>2 in *Mll3/4* DKO versus WT mESCs. **(C)** Heatmaps centered either on 10,462 DNTTIP1 downregulated peaks (FC>2) (upper) or on 5,513 regions with decreased H4K20ac (FC>2) (lower) in *Mll3/4* DKO compared to WT mESCs. Occupancy of DNTTIP1, H4K20ac, H3K4me1, H3K4me2, H3K4me3, H3K27ac, and H3K27me3 in WT and *Mll3/4* DKO mESCs is displayed. **(D)** Genome browser tracks depicting the ChIP-seq profiles of DNTTIP1, H4K20ac, H3K4me1, H3K4me2, H3K4me3, H3K27ac, and H3K27me3 in WT and *Dnttip1* KO mESCs, and WT and *Mll3/4* DKO mESCs at the promoter and two enhancer regions of the *Spry4* locus.

**Figure 5.**
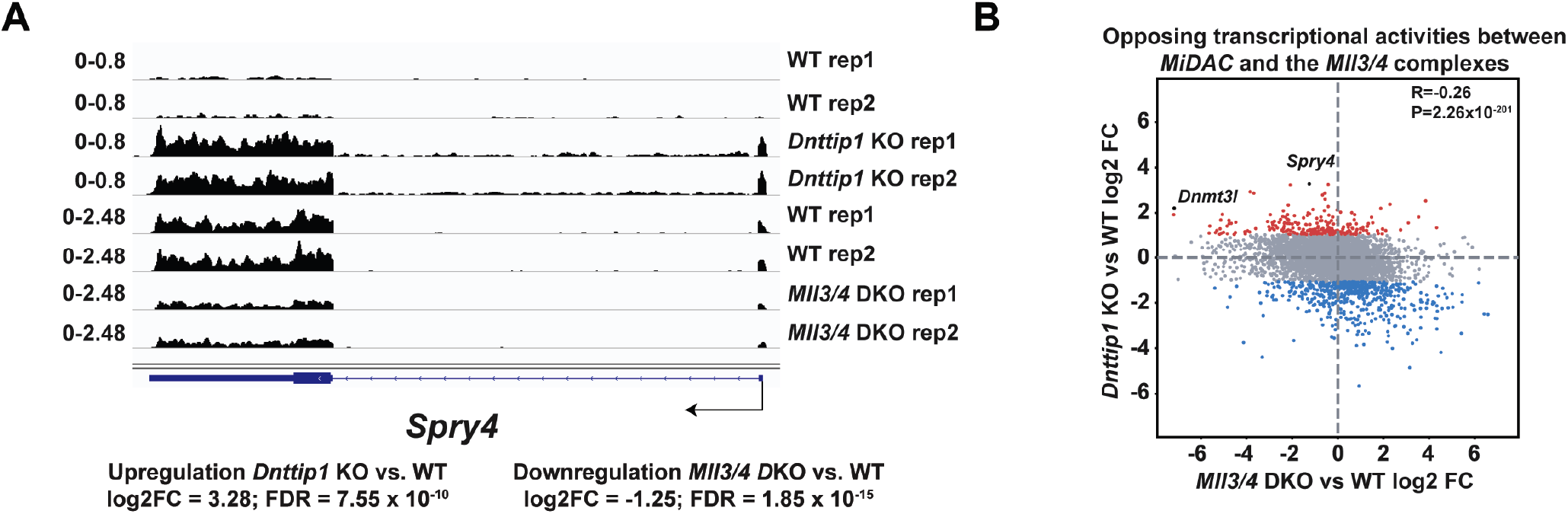
A specific gene expression program is controlled by the opposing functions of MiDAC and the MLL3/4 complexes. **(A)** Genome browser tracks showing the RNA-seq profile of the *Spry4* gene in WT and *Dnttip1* KO mESCs, and WT and *Mll3/4* DKO mESCs. Two replicates are displayed for each genotype. **(B)** Scatter plot depicting the log_2_ fold change of all expressed genes in *Dnttip1* KO versus WT mESCs (y-axis) and *Mll3/4* DKO versus WT mESCs (x-axis). A significant anticorrelation of differentially expressed genes that are transcriptionally controlled by both MiDAC and the MLL3/4 complexes is observed.

## Discussion

MiDAC is the least studied class I histone deacetylase complex and its important physiological role in neurogenesis and embryonic development has only very recently been described (Mondal et al 2020, Turnbull et al 2020). How MiDAC functionally intersects with other epigenetic regulators to elicit changes in chromatin state and effect transcriptional responses is currently unknown. Here, we report for the first time a functional link between MiDAC and the MLL3/4 complexes. We show that UTX, a complex-specific subunit of the MLL3/4 complexes, acts as an important hub to ensure the physical association with MiDAC via its scaffolding subunit ELMSAN1 (Figure 1). Indeed, apart from mediating the interaction with MiDAC, UTX might generally function as a unique docking site within the MLL3/4 complexes to recruit other chromatin-associated protein complexes, including the previously reported TOP (<u>T</u>ET2, <u>O</u>GT, and <u>P</u>ROSER1) complex (Wang et al., 2022). Interestingly, *UTX* is often mutated in the same cancer types as *MLL3* and/or *MLL4* linking the function of the MLL3/4 complexes directly to carcinogenesis via UTX (Martinez-Jimenez et al 2020). Thus, gaining mechanistic insight into the role of UTX-interacting chromatin-modifying complexes such as MiDAC and the TOP complex might open an avenue to specifically target these cancers. The results of our study implicate MiDAC and the MLL3/4 complexes as antagonistic co-regulators of a specific gene expression program (Figure 6). This relationship is clearly evident at the level of histone modifications. We previously identified MiDAC as a major negative regulator of H4K20ac (Mondal et al 2020) and here provide evidence that this function of MiDAC towards H4K20ac extends to many gene regulatory elements genome-wide (Figure 2). Conversely, the MLL3/4 complexes function to promote H4K20ac deposition on many loci that are co-regulated by the MLL3/4 complexes and MiDAC (Figure 4). These findings imply that the degree of H4K20ac enrichment on promoters and putative enhancers might be used as an indicator to predict the transcriptional response of a gene expression program that is balanced by the opposing action of MiDAC and the MLL3/4 complexes. To date our understanding of H4K20ac is extremely limited. Thus, future work is required to further elucidate the role of H4K20ac in transcription and potentially other processes. One possible mechanism by which the MLL3/4 complexes might regulate H4K20ac could be through direct binding to acetylated histones. For example, previous studies have shown that the PHD6 domain of MLL4 is able to recognize and bind H4K16ac (Zhang et al 2019). Therefore, it is possible that other PHD domains or regions within MLL3 and/or MLL4 or domains/regions within UTX might recognize H4K20ac and thus protect H4K20ac from being deacetylated by MiDAC. Alternatively, the MLL3/4 complexes might promote the activity of a histone acetyltransferase that is able to target H4K20ac.

**Figure 6.**
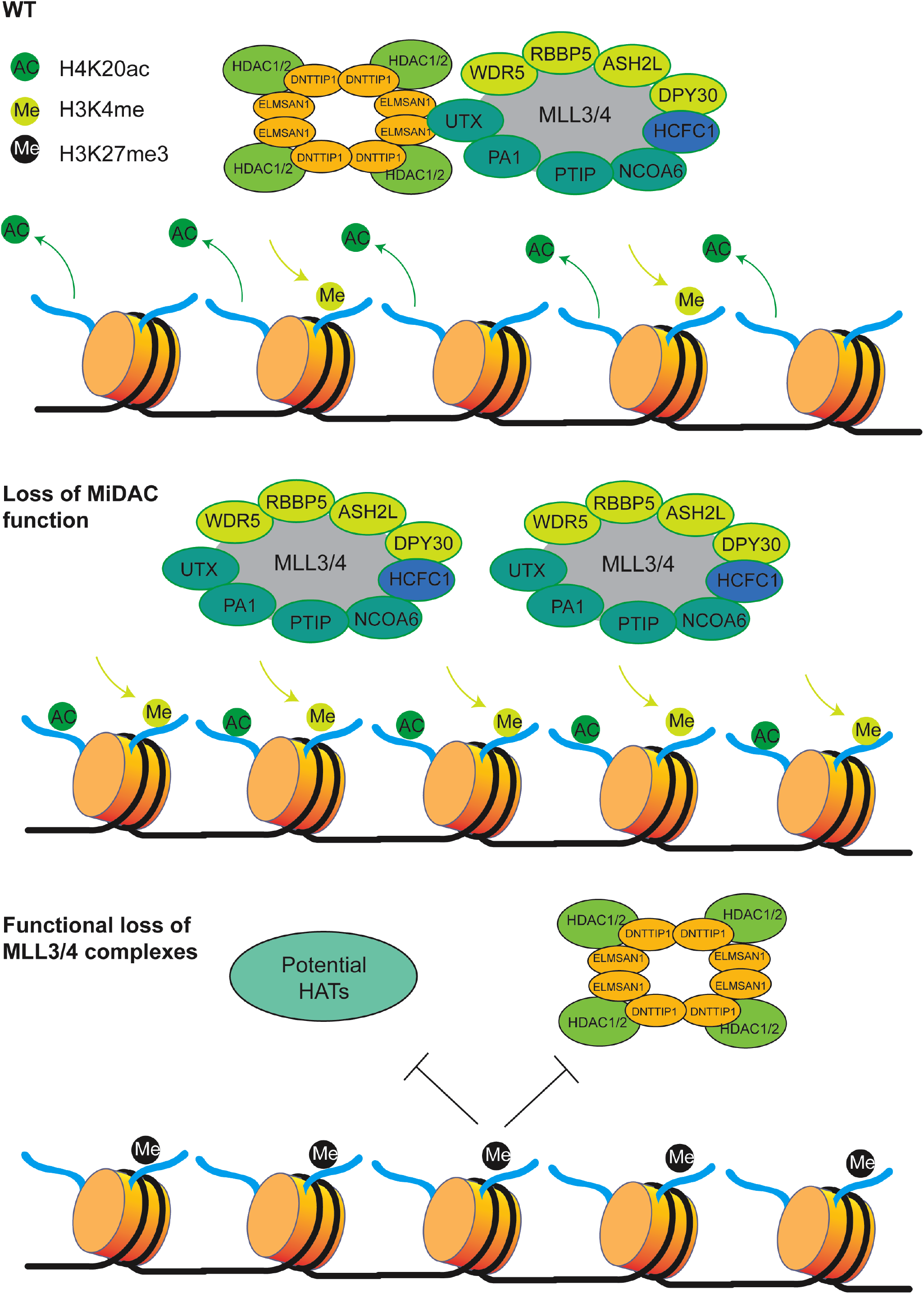
Model describing the interplay between MiDAC and MLL3/4 complexes. Under WT conditions, MiDAC functionally intersects with the MLL3/4 complexes through the interaction between ELMSAN1 and UTX. The removal of H4K20ac and deposition of H3K4me2 are in a balanced state. Upon loss of MiDAC, H4K20ac increases genome-wide and by an unknown mechanism the MLL3/4 complexes’ deposition also increases, thus increased H3K4me2 on genomic elements that are co-regulated by MiDAC and the MLL3/4 complexes. This indicates that under normal conditions, MiDAC may play an inhibitory role on MLL3/4 complexes. Upon loss of the MLL3/4 complexes, MiDAC is reduced genome-wide which intriguingly does not cause the increase of H4K20ac the substrate of MiDAC, which may be partially due to elevated H3K27me3 preventing the access of potential histone acetyltransferases (HATs) and MiDAC.

Unexpectedly, we found that the decrease of H4K20ac on gene regulatory elements in *Mll3/4* DKO mESCs did not coincide with higher MiDAC occupancy on these loci but rather a decrease in MiDAC enrichment. As the protein levels of MiDAC components are unchanged in *Mll3/4* DKO mESCs (Figure S6), this suggests that the genomic localization of MiDAC is dependent on the MLL3/4 complexes and that in the absence of the MLL3/4 complexes the inhibitory role of MiDAC toward H4K20ac is suppressed. It is possible that the MLL3/4 complexes are directly involved in the recruitment of MiDAC to its target loci or that an unknown MiDAC recruiting factor is reduced in *Mll3/4* DKO mESCs. The strong decrease of H3K4me1 in *Mll3/4* DKO mESCs could also cause the reduction in MiDAC occupancy, as MiDAC deposition may be dependent on direct binding to H3K4me1. Interestingly however, we observed higher enrichment of the repressive histone mark H3K27me3 at loci with reduced MiDAC occupancy or decreased H4K20ac enrichment in *Mll3/4* DKO mESCs. Thus, the formation of a repressive chromatin state (via H3K27me3) may prevent MiDAC and potential histone acetyltransferases that target H4K20 to access those regions. Furthermore, we cannot exclude that one or multiple H4K20 deacetylase(s) is/are activated and/or target H4K20ac more effectively in *Mll3/4* DKO mESCs. A potential model of the interplay between MiDAC and the MLL3/4 complexes is described in Figure 6. Besides H4K20ac, other histone marks such as H3K4me2 and to some degree H3K27ac are also antagonistically regulated by MiDAC and the MLL3/4 complexes (Figure 2-4). It has previously been shown that MLL3/4 can catalyze both H3K4me1 and H3K4me2 on enhancers (Hu et al 2013, Lee et al 2013). However, the increased occupancy of UTX and MLL4 we observed in *Dnttip1* KO mESCs on regulatory elements that are also targeted by MiDAC did not result in higher H3K4me1 enrichment, but selectively appeared to be confined to increased H3K4me2 and H3K27ac enrichment (Figure 2 and 3). One potential reason why we observed no significant changes of H3K4me1 in *Dnttip1* KO mESCs may be due to an increased propensity of the MLL3/4 complexes to catalyze H3K4me2 rather than H3K4me1 in the presence of H4K20ac. Finally, our transcriptome-wide analysis clearly confirms the antagonistic relationship between MiDAC and the MLL3/4 complexes and aligns well with our observations on the level of histone modifications as discussed above (Figure 5). Future studies will be required to further interrogate the role of MiDAC and the MLL3/4 complexes in development and disease.

## Materials and Methods

### Cell lines

*Dnttip1* KO (clone 1-D1E6) and their wild-type control mESCs were previously described by our lab (Mondal et al., 2020). *Mll3/4* DKO, *Mll3/4* DCD and their wild-type control mESCs were kindly provided by the Wysocka lab (Dorighi et al., 2017). All mESCs were cultured under chemically defined naïve culture conditions (2iL) in 0.1% gelatin-coated flasks or plates. *ELMSAN1 TRERF1* DKO HEK293 cells were generated from wild-type Flp-In T-REx HEK293 cells (Invitrogen, R78007) by the Center for Advanced Genome Engineering (CAGE) at St. Jude Children’s Research Hospital using CRISPR/Cas9-mediated gene editing (Figure S2). HCT116 cells (*MLL3-/- MLL4+/+*) were obtained from ATCC. *MLL3*+/+ *MLL4*+/+ HCT116 cells and *MLL3*-/- *MLL4-*/- HCT116 cells were generated from *MLL3*-/- *MLL4*+/+ HCT116 cells by the CAGE at St. Jude Children’s Research Hospital using CRISPR/Cas9-mediated gene editing (Figure S3A). Isogenic tetracycline-inducible FLAG-UTX and FLAG control HEK293 cells were previously described by our lab (Wang et al 2022). Isogenic tetracycline-inducible FLAG- DNTTIP1, FLAG-ELMSAN1, FLAG-TRERF1 and FLAG control HEK293 cells were generated by using Flp recombinase-mediated integration in *ELMSAN1 TRERF1* DKO HEK293 cells. HEK293 and HCT116 cells were cultured in Dulbecco’s Modified Eagle Medium (Gibco, 11995065) with 10% FBS (VWR, 97068-085) and 1% Penicillin/Streptomycin (Gibco, 15140122). All cell lines tested negative for mycoplasma contamination.

### Antibodies

#### For Western Blotting

Mouse α-Actin (Developmental Studies Hybridoma Bank [DSHB], JLA20 supernatant) at 1:1,000; rabbit α-ASH2L (Cell Signaling Technology, 5019S) at 1:2,000; rabbit α-DNTTIP1 (Bethyl Laboratories, A304-048A) at 1:2,000; rabbit α-ELMSAN1 (Bethyl Laboratories, A303-157A) at 1:5,000; rabbit α-ELMSAN1 (Herz lab, 34421) at 1:5,000; mouse α-FLAG (Sigma, F3165) at 1:2,000; rabbit α-HDAC1 (Cell Signaling Technology, 34589S) at 1:2,000; rabbit α-HDAC2 (Cell Signaling Technology, 57156S) at 1:2,000; rabbit α-KAT2A (Cell Signaling Technology, 3305S) at 1:2,000; rabbit α-MLL3/KMT2C (Herz lab, #31865 and #31866 (both human aa 581-850) at 1:5,000; rabbit α-MLL4/KMT2D (Herz lab, #31863 (human aa 1-181) and #32757 (human aa 281-506) at 1:5,000; rabbit α-RBBP5 (Cell Signaling Technology, 13171S) at 1:2,000; rabbit α-TRERF1 (Sigma, HPA051273) at 1:2,000; rabbit α-UTX/KDM6A (Cell Signaling Technology, 33510S) at 1:2,000; α-WDR5 (Cell Signaling Technology, 13105S) at 1:2,000.

#### For ChIP-seq

Rabbit α-H3K4me1 (RevMAb, 31-1046-00), 10 µg per ChIP; mouse α-H3K4me2 (Active Motif, 39679), 10 µg per ChIP; rabbit α-H3K4me3 (RevMAb, 31-1039-00), 10 µg per ChIP; rabbit α- H3K27ac (RevMAb, 31-1056-00), 10 µg per ChIP; rabbit α-H3K27me3 (Cell Signaling Technology, 9733S), 30 µl per ChIP; rabbit α-H4K20ac (RevMAb, 31-1084-00), 10 µg per ChIP; rabbit α-DNTTIP1 (Bethyl Laboratories, A304-048A), 10 µg per ChIP; rabbit α-UTX/KDM6A (Cell Signaling Technology, 33510S), 30 µl per ChIP; rabbit α-MLL4/KMT2D (Ge lab, #3), 10 µg per ChIP.

### CRISPR/Cas9 gene editing

*ELMSAN1 TRERF1* DKO clones from Flp-In T-REx HEK293 and *MLL3*+/+ *MLL4*+/+ clones and *MLL3*-/- *MLL4-*/- clones from HCT116 cells were generated using CRISPR/Cas9 technology. Briefly, 400000 Flp-In T-REx HEK293 cells (Thermo Fisher Scientific, R78007) were transiently transfected with precomplexed ribonuclear proteins (RNPs) consisting of 100 pmol of each chemically modified sgRNA (Synthego), 70 pmol of Cas9 protein (St. Jude Protein Production Core), and 200 ng of pMaxGFP (Lonza) via nucleofection (Lonza, 4D-Nucleofector™ X-unit) using solution P3 and program CM130 in a small (20 µl) cuvette according to the manufacturer’s recommended protocol. Five days post transfection, cells were single cell sorted by FACS to enrich for GFP-positive (transfected) cells, clonally selected and verified for the desired targeted modification via targeted deep sequencing. Targeted amplicons were generated using gene specific primers with partial Illumina adapter overhangs and sequenced as previously described (Sentmanat et al 2018). Briefly, cell pellets of approximately 10000 cells were lysed and used to generate gene specific amplicons with partial Illumina adapters in PCR #1. Amplicons were indexed in PCR #2 and pooled. Additionally, 10% PhiX Sequencing Control V3 (Illumina) was added to the pooled amplicon library prior to running the sample on a Miseq Sequencer System (Illumina) to generate paired 2 x 250 bp reads. Samples were demultiplexed using the index sequences, fastq files were generated, and NGS analysis was performed using CRIS.py (Connelly & Pruett-Miller 2019). Two clones were initially identified, and one was used for further characterization as it pertains to this manuscript. Editing construct sequences and relevant primers are listed in Table S2.

### Immunoprecipitation (IP)

Large scale IPs from ten 150 mm plates and small scale IPs from one 150 mm plate were carried out as previously reported (Wang et al 2022).

### Glycerol gradient fractionation

Glycerol gradient fractionation was conducted as previously described (Wang et al 2022). 34 fractions of approximately 325 μl each were collected and analyzed by western blotting for UTX, MLL3, MLL4, RBBP5, ELMSAN1, DNTTIP1, HDAC1 and HDAC2.

### ChIP-seq

ChIP-seq was performed as previously described (Mondal et al 2020, Wang et al 2022).

### Mass spectrometry (MS)

#### Protein identification by liquid chromatography coupled with tandem mass spectrometry

##### Sample preparation

Protein samples were briefly run into a 4-20% PAGE gradient gel as described in a previously published protocol (Xu et al 2009). The gel bands were destained, reduced with dithiothreitol (DTT), alkylated by iodoacetamide (IAA), washed, dried down, and rehydrated with a buffer containing trypsin. After overnight proteolysis at 37°C, peptides were extracted, dried down in a speed vacuum and reconstituted in 5% formic acid.

##### Mass spectrometry

The peptide mixture from each gel band was separated on a nanoscale capillary reverse phase C18 column (75 µm id, 10 cm) by a HPLC system (Thermo EASY-nLC 1000). Buffer A was 0.2% formic acid and Buffer B was 0.2% formic acid in 70% acetonitrile. The peptides were eluted by increasing Buffer B from 12% to 70% over a 60-90 min gradient. The peptides were ionized by electrospray ionization and detected by a Thermo LTQ Orbitrap Elite mass spectrometer. The mass spectrometer was operated in data-dependent mode. For each duty cycle a high resolution survey scan in the Orbitrap and 20 low resolution MS/MS scans were acquired in the ion trap.

##### Database search and analysis

The MS data were searched against the human UniProt database using Sequest (version 28, rev. 12) (Eng et al 1994). The database was concatenated with a reversed decoy database for evaluating false discovery rate (Peng et al 2003). Mass tolerance of 15 ppm for precursor ions and 0.5 Da for product ions were used. Two missed cleavages with a maximum of three modifications were allowed and assignment of b, and y ions were used for identification. Carbamidomethylation of Cysteine (+57.02146 Da) for static modification and oxidation of Methionine (+15.99492 Da) for dynamic modification were considered. Mass accuracy and matching scores filters were used for MS/MS spectra to reduce the protein false discovery rate to < 1%. Spectral counts of each protein may reflect their relative abundance in the samples after normalizing for protein molecular weight. The spectral counts between the samples for a given protein were used to calculate the p-value which was derived by G-test (Bai et al 2013).

#### Relative quantification of histone post translational modification abundances using LC-MS/MS

##### Sample preparation

Histones were acid extracted as described previously (Shechter et al 2007). In brief, mESCs were lysed in 10X cell pellet volume of ice-cold hypotonic lysis buffer (15 mM Tris-HCl (pH 7.5), 60 mM KCl, 11 mM CaCl_2_, 5 mM NaCl, 5 mM MgCl_2_, 250 mM sucrose, 1 mM dithiothreitol, 10 mM sodium butyrate) supplemented with 0.1% NP-40 on ice for 5 min. Nuclei were pelleted by centrifugation (1,000g, 2 min, 4^○^C) and washed twice in ice-cold hypotonic lysis buffer w/o NP-40. Nuclei were resuspended in 5X nuclei pellet volumes of ice-cold 0.2 M sulfuric acid and mixed on a rotation wheel for 120 min at 4^○^C. Insolubilized nuclear debris was pelleted by centrifugation (16,000g, 10 min, 4^○^C). Supernatant was transferred to a fresh low-protein binding Eppendorf tube and histone proteins were precipitated by adding ice-cold trichloroacetic acid (TCA) to the final concentration of 20% (v/v) followed by 60 min incubation on ice. Precipitated histone proteins were pelleted by centrifugation (16,000g, 10 min, 4^○^C), washed 3 times with acetone (-20^○^C) and resuspended in MS grade water.

##### Mass spectrometry

Extracted histones were prepared for LC-MS/MS analysis using the hybrid chemical derivatization method as described previously (Maile et al 2015). In brief, 4 μg aliquots of purified histones were diluted with MS grade water to a total volume of 18 μl and buffered to pH 8.5 by addition of 2 μl of 1 M triethylammonium bicarbonate buffer (TEAB). Propionic anhydride was mixed with MS grade water in a ratio of 1:100 and 2 μl of the anhydride-mixture was added immediately to the histone sample, with vortexing, and the resulting mixture was incubated for 5 min at room temperature. The reaction was quenched by adding 2 μl of 80 mm hydroxylamine followed by 20 min incubation at room temperature. Tryptic digestion was performed overnight with 0.5 μg trypsin per sample at 37^○^C. A 1% v/v solution of phenyl isocyanate (PIC) in acetonitrile was freshly prepared and 6 μl added to each sample and incubated for 60 min at 37 °C. Samples were acidified by adding trifluoroacetic acid (TFA) to the final concentration of 1%. Peptides were de- salted with C18 spin columns (Pierce ™) following the manufacture protocol. Peptides were eluted from C18 spin columns with 70% acetonitrile, partially dried in a speedvac and resuspended in 30 μl 0.1% TFA.

The resulting peptide mixtures were analyzed using nano-flow liquid chromatography tandem mass spectrometry (LC-MS/MS) on a Q-Exactive HF mass spectrometer coupled to an Ultimate 3000 nano- UPLC (Ultimate 3000, Dionex, Sunnyvale, CA) in data-dependent acquisition (DDA) mode. ∼200 ng peptide aliquot was used per one sample per one injection. Peptides were loaded automatically on a trap column (300 µm inner diameter ×5 mm, Acclaim PepMap100 C18, 5 µm, 100 Å; LC Packings, Sunnyvale, USA) prior to C18 reversed phase chromatography on the analytical column (nanoEase MZ HSS T3 Column, 100 Å, 1.8 µm, 75 µm × 250 mm; Waters, Milford, USA). Peptides were separated at flowrate of 0.250 μl per minute by a linear gradient from 1% buffer B (0.1% (v/v) formic acid, 98% (v/v) acetonitrile) to 25% buffer B over 40 min followed by a linear gradient to 40% B in 20 min, then to 85% B in 5 min. After 5 min at 85% buffer B, the gradient was reduced to 1% buffer B over 2 min and then allowed to equilibrate for 8 min. Full mass range spectra were at 60,000 resolution (at *m*/*z* 400), and product ions spectra were collected in a “top 15” data dependent scan cycle at 15,000 resolution.

##### Data analysis

RAW MS data were analyzed using EpiProfile 2.0 software (Yuan et al 2018). The reported relative abundances of histone modifications were validated manually using an open-source Skyline software.

### ChIP-seq data processing

Following the procedure described before (Wang et al., 2022), raw sequencing reads were pre- processed with the Trim-Galore tool (v0.4.4, https://www.bioinformatics.babraham.ac.uk/projects/trim_galore/) (Krueger et al 2012) and cutadapt (DOI:10.14806/ej.17.1.200), to remove low quality reads, remove potential adapters and quality trim reads’ 3′ ends. Quality score cutoff was set to Q20. Next, the remaining reads were mapped to the mouse reference genome (mm10) with bwa aln, followed by bwa samse (Li & Durbin 2009) (v0.7.12-r1039) with -K flag set to 10,000,000. The output was then converted to binary alignment map (BAM) format with samtools (Li et al 2009) (v1.2). Next, the bamsormadup tool from biobambam2 (v2.0.87, DOI: 10.1186/1751-0473-9-13) was used to identify duplicated reads, and the SPP tool (Kharchenko et al 2008) (v1.11), was used to conduct the Cross-Correlation analysis and estimate the fragment size. Subsequently, samtools was used again in order to extract uniquely mapped reads, and bedtools (Quinlan & Hall 2010) (v2.24.0) was then used to extend the reads with the previously estimated fragment size. The intermediate files containing the extended fragments, were converted to bigwig track files by University of California, Santa Cruz (UCSC) tools (Kuhn et al 2013) (v4), and signal intensity was corrected for sequencing depth, normalizing them to 15 million uniquely mapped non-duplicated fragments. For visualization purposes, for the experiments that per condition consisted of more than one replicate, the bigwig files from individual replicates were merged calculating average per bin signal between replicates. Subsequently, MACS2 program (Zhang et al 2008) was used to call peaks in narrow mode, with - -nomodel -q 0.05 flags (high confidence peaks). Separately, narrow peaks were also called with more relaxed criteria, setting the -q flag to 0.5, which are here referred to as FDR50 peaks. Next, for experiments with more than one replicate, reproducible peaks (e.g. for H4K20ac in *Dnttip1* KO mESCs) were identified as those with overlapping FDR50 peaks present in all replicates at a given genomic region. Otherwise, for ChIP-seq targets without replicates, only the high confidence peaks were considered. Finally, the reproducible peaks from the same immunoprecipitation target (e.g. H4K20ac) were merged into the collection of reference peaks, here further referred to as all reproducible peaks.

### Differential binding peak identification

To perform statistical testing between experimental groups, the number of fragments for each reference peak was counted with intersect command from pybedtools (Dale et al 2011, Quinlan & Hall 2010) (v0.8.1). Next, the number of raw fragments mapping per peak was converted to FPKM units (Fragments Per Kilo base per Million mapped reads); and then TMM (trimmed mean of M-values) from edgeR (Robinson et al 2010), followed by the limma-voom approach (Ritchie et al 2015) was used to assess the significance of differential peak binding. For the contrasts, for which based on previously published independent work (Dorighi et al 2017, Mondal et al 2020), a genome-wide change (either gain or loss) was expected, and for which no spike-in was available, an additional pre-processing step was introduced to calculate the scaling factors. The contrasts for this step included DNTTIP1, H4K20ac, UTX, and MLL4 in *Dnttip1* KO mESCs and H4K20ac, H3K4me1 and DNTTIP1 in *Mll3/4* DKO mESCs. Those scaling factors were computed by first calculating the median of single base-pair resolution enrichment signal from the bigwig track files, corrected previously for sequencing depth (see above), which was accomplished with the pybigwig tool (available online at: https://github.com/deeptools/pyBigWig), over all reproducible peaks. Next, the median signal from each sample was multiplied by 1/1,000,000, and converted to integer, which value imitated the spike-in read counts for the purpose of the scaling factor calculation and is further referred to as pseudo-spike reads. Next, similar to the approach used by the authors of DESeq2 and edgeR (Love et al 2014, Robinson et al 2010) the scaling factors were calculated normalizing individual pseudo-spike reads to the maximum pseudo-spike reads count across samples, and then dividing these values by their geometric mean. Scaling factors calculated this way, were further supplied to the *norm.factors* parameter of the DGElist function of edgeR for differential peak calling. For the contrasts with replicates, the region was considered as differentially binding, when the false discovery rate (FDR) was lower than 0.05 and the log_2_(fold-change) > 1, for increased binding, or log_2_(fold-change) < -1 for decreased binding. For some experiments without available biological replicates (including UTX and MLL4 for *Dnttip1* KO mESCs and H3K4me1, H3K4me3 and H3K27Ac for *Mll3/4* DKO mESCs), the identification of differentially bound regions was based on the fold-change value, using a log_2_(fold-change) > 1 threshold to identify increased binding and log_2_(fold-change) < -1 for decreased binding.

### Annotation of genomic regions

Following the approach used previously (Wang et al., 2022), genomic regions were assigned to their genomic contexts with an in-house script based on pybedtools (Dale et al 2011) (v0.8.1), such that each region could only be assigned to one feature. For this purpose, genomic regions were successively overlapped with predefined genomic contexts in the following prioritization order: (1) Promoter.Up: region up to 2 kbp upstream from TSS; (2) Promoter.Down: region up to 2 kbp downstream from TSS; (3) Exons; (4) Introns; (5) TES: transcription end sites; (6) 5′ Distal: region up to 50 kbp upstream from TSS, excluding promoter region; (7) 3′ Distal: region up to 50 kbp downstream from TSS, excluding promoter region; (8) Intergenic. The reference annotation for TSS, and all subsequent genomic contexts, was based on the Gencode vM14 (Frankish et al 2019) reference annotation and included all isoforms. In parallel, the genomic regions were annotated with genes, via putative promoter-related association. For that purpose, genomic regions were overlapped with promoter regions with bedtools (Quinlan & Hall 2010) (v2.24.0); one region could be assigned to multiple genes. The promoter region was defined as TSS ± 2kbp. Next, regions not assigned to any gene as promoter-associated, were assigned to a gene as putative enhancer-related regions, if their distance to the gene’s TSS was within a threshold of ± 50 kbp, excluding the promoter region.

### RNA-seq data processing

Sequenced RNA-seq reads were quality-filtered using TrimGalore (https://www.bioinformatics.babraham.ac.uk/projects/trim_galore/), and then aligned to the mouse reference genome (mm10) using STAR (Dobin et al 2013). Next, RSEM was used to quantify read counts per gene (Dobin et al 2013). Subsequently, differential expressed genes were identified using the limma-voom approach (Law et al 2014, Ritchie et al 2015), as previously described (Wang et al 2022).

### Gene set enrichment analysis

Gene set enrichment analysis (GSEA) was conducted with GSEApy (Dale et al 2011, Mootha et al 2003) (v0.10.4, available on-line at https://gseapy.readthedocs.io/en/latest/), using the pre-ranked list of genes, where the per gene metric was an equivalent of the log_2_(fold-change) values, derived from differential gene expression analyses. GSEApy’s prerank function was used with standard parameters, except –min-size and –max-size flags, whose values were set to 5 and 5,000, respectively. The GSEA analysis was run with the collection of gene sets from the KEGG database (Kanehisa & Goto 2000), downloaded from the Enrichr portal (Kuleshov et al 2016). The above-mentioned gene sets collection was additionally expanded with in-house gene sets, representing genes annotated with various genomic regions, e.g. regions with increased H4K20ac, or genomic regions displaying both an increase in UTX and MLL4 occupancy. Following previous approaches (Subramanian et al 2005) (DOI: 10.1007/978-3-540-73731-5_2) we considered the gene sets that are significantly associated with positive or negative phenotype as the ones whose FDR was lower than 25% and whose p-value was lower than 0.05.

### Data Availability

RNA-seq data from WT and *Dnttip1* KO mESCs was obtained from GSE131062. RNA-seq data from WT and *Mll3/4* DKO mESCs was obtained from GSE98063. DNTTIP1 ChIP-seq data in WT and *Dnttip1* KO mESCs was used from GSE131062. The accession number for the remaining ChIP-seq datasets reported in this manuscript is GEO: GSE190323.

## Acknowledgements

We thank the Wysocka lab for providing *Mll3/4* DKO mESCs and their WT controls. We gratefully acknowledge the following research resources at St. Jude Children’s Research Hospital for their services: the Hartwell Center for ChIP-seq library preparation and sequencing, the center for Proteomics and Metabolomics for conducting mass spectrometry analysis, and Center for Applied Bioinformatics for bioinformatic analysis. The mouse α-Actin monoclonal hybridoma antibody (JLA20) developed by the University of Iowa was obtained from the Developmental Studies Hybridoma Bank, created by the NICHD of the NIH and maintained at The University of Iowa, Department of Biology, Iowa City, IA 52242. We thank everyone in the Herz lab for their helpful comments and critical reading of the manuscript. This work was funded by a transition to independence grant from the National Institutes of Health/National Cancer Institute (R00CA181506 to H-MH, P30 CA021765 to SMP-M); and the American Lebanese Syrian Associated Charities (ALSAC). The content is solely the responsibility of the authors and does not necessarily represent the official views of the National Institutes of Health.

## Author Contributions

H-MH and XW conceived and designed the study. H-MH, XW, YS, BM and AT performed the experiments and analyzed the data. WR and BS performed computational analysis. RB and SMPM generated knockout cells. H-MH and XW wrote the manuscript. SMPM, RS and H-MH supervised the study.

## Competing Interests

All authors declare no competing interests.

## Supplementary Figure Legends

**Figure S1.**
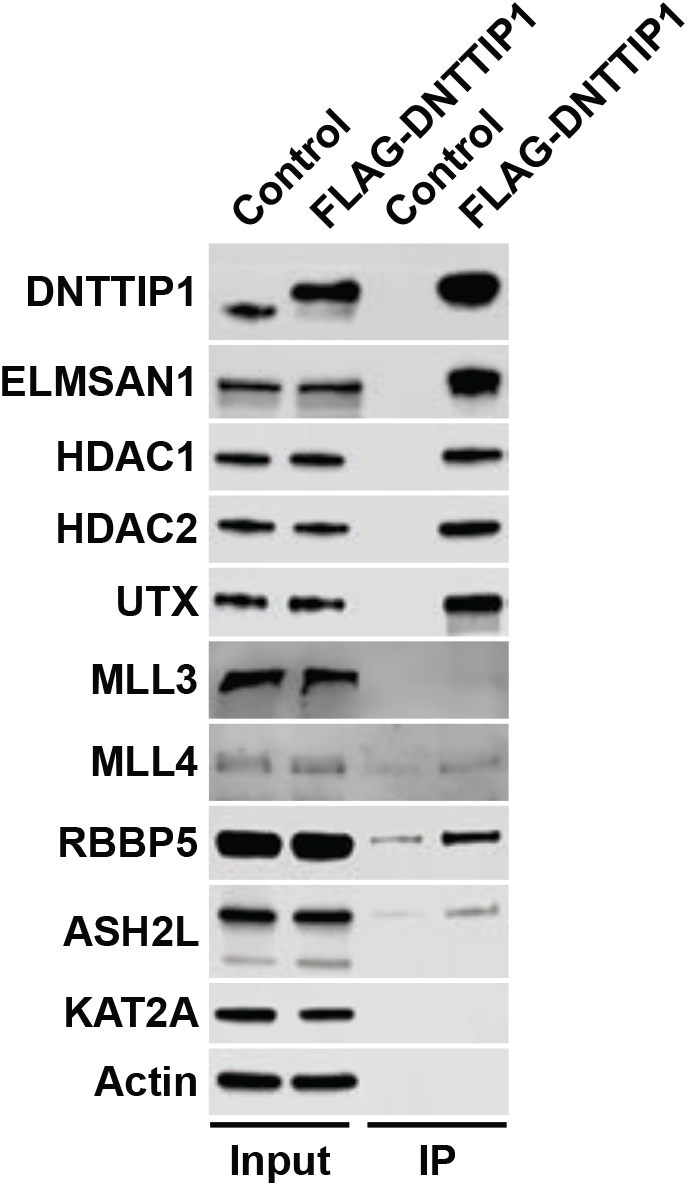
The MLL3/4 complexes associate with the mitotic deacetylase complex (MiDAC). Western blot (WB) of FLAG-DNTTIP IP from HEK293 cells confirming interaction of DNTTIP1 with the MiDAC components ELMSAN1, HDAC1, HDAC2 and members of the MLL3/4 complexes including UTX, the H3K4 methyltransferases MLL3 and MLL4, and the two core components RBBP5 and ASH2L. KAT2A was used as a negative control. HEK293 cells with a FLAG-tag expressing plasmid were used as an IP control. Nuclear extracts were used as input. Actin was used as a loading control for the inputs.

**Figure S2.**
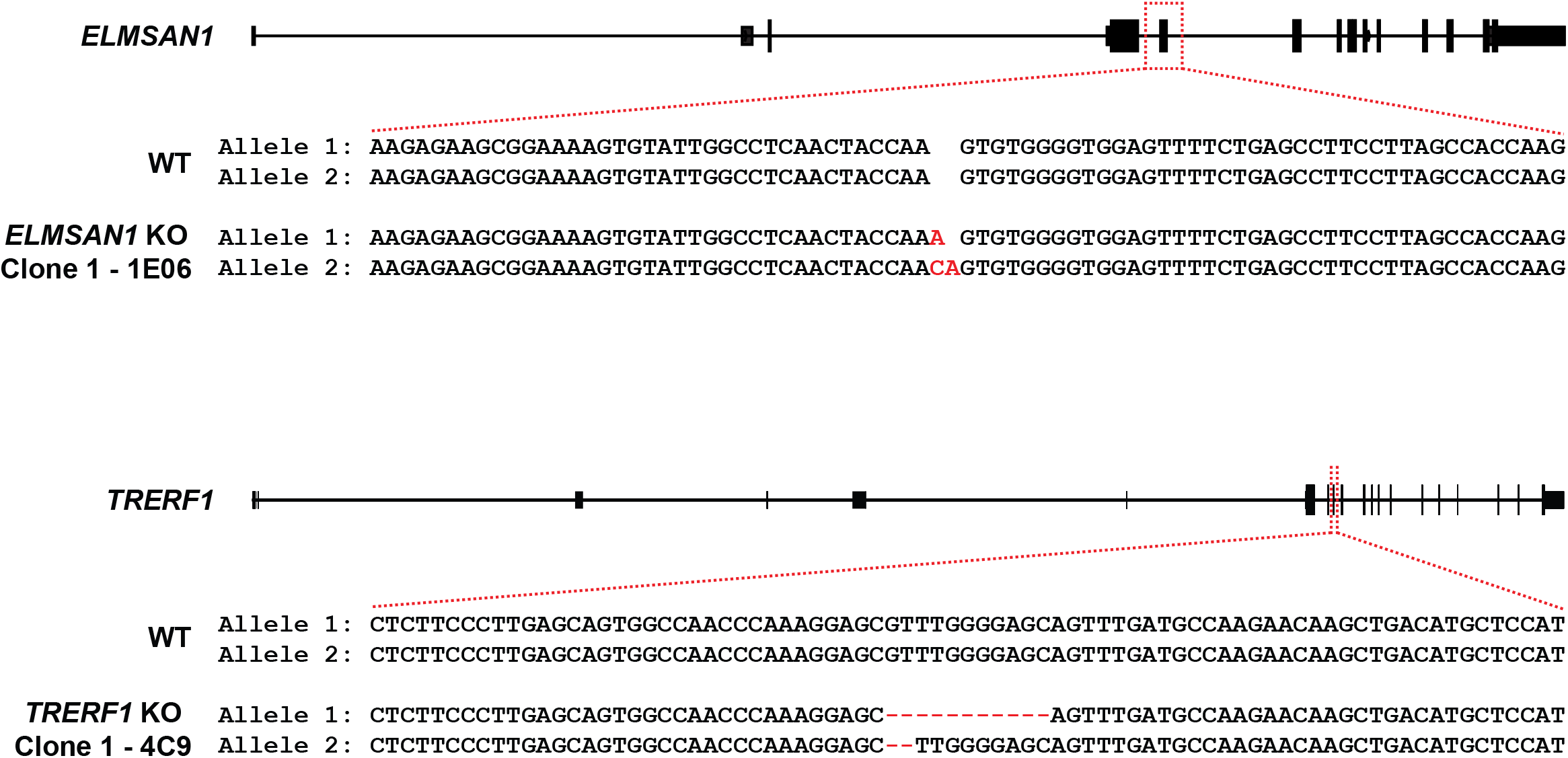
CRISPR KO strategy to create *ELMSAN1 TRERF1* DKO clones in HEK293 cells. The *ELMSAN1* locus was targeted with gRNA SM109.hELMSAN1.g1 in Flp-In T-Rex HEK293 cells to create *ELMSAN1* KO clone 1 – 1E06. The *TRERF1* locus was targeted with gRNA CAGE654.TRERF1.g2 in *ELMSAN1* KO clone 1 to create *ELMSAN1 TRERF1* DKO clone 1 – 4C9. The created indels for allele 1 and 2 of *ELMSAN1* in *ELMSAN1* KO clone 1 and for allele 1 and 2 of *TRERF1* in *ELMSAN1 TRERF1* DKO clone 1 are highlighted in red.

**Figure S3.**
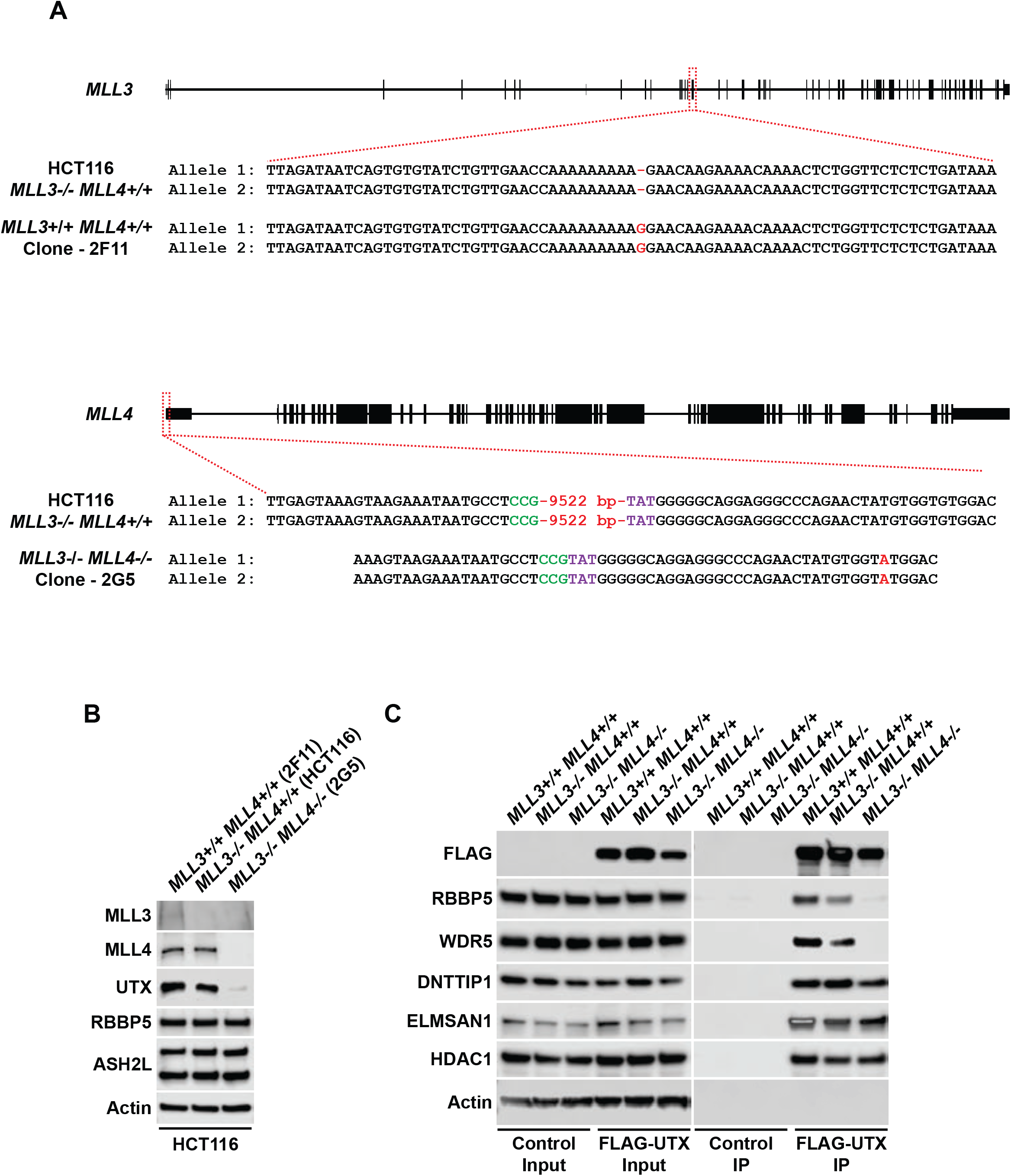
The interaction interface between UTX and ELMSAN1 forms the nexus between the MLL3/4 complexes and MiDAC. **(A)** CRISPR strategy to restore MLL3 expression and knock out MLL4 in HCT116 cells. A single base pair deletion in the *MLL3* locus was repaired by utilizing a single-stranded oligo donor that contained a blocking modification (G>A mutation at the highlighted position) to create *MLL3+/+ MLL4-/-* clone 2F11. The newly added G to restore the *MLL3* sequence back to WT in HCT116 cells is highlighted in red. The *MLL4* promoter was targeted with gRNAs SM124.KMT2D.g4 and SM125.KMT2D.g3 resulting in the deletion of a 9522 bp sequence within the *MLL4* promoter to create *MLL3-/- MLL4-/-* clone 2G5. The bases at the deletion junction are marked by green and purple, respectively. Additionally, a G to A substitution in allele 1 and 2 (highlighted in red) was created 33 bp from the deletion junction. **(B)** WB for members of the MLL3/4 complexes including UTX, the H3K4 methyltransferases MLL3 and MLL4 and the core components RBBP5 and ASH2L from nuclear extracts of *MLL3+/+ MLL4+/+* (2F11), *MLL3-/- MLL4+/+* (control) and *MLL3-/- MLL4-/-* (2G5) HCT116 cells. Actin was used as a loading control. **(C)** WB of FLAG-UTX IP from *MLL3+/+ MLL4+/+* (2F11), *MLL3-/- MLL4+/+* (control) and *MLL3-/- MLL4-/-* (2G5) HCT116 cells after transfection with FLAG-UTX. UTX is able to associate with MiDAC (DNTTIP1, ELMSAN1 and HDAC1) in the absence of MLL3 and MLL4 suggesting that UTX mediates the interaction between the MLL3/4 complexes and MiDAC. HCT116 cells of the same genotypes transfected with a FLAG-tag expressing plasmid were used as IP controls. Nuclear extracts were used as input. Actin was used as a loading control for the inputs.

**Figure S4.**
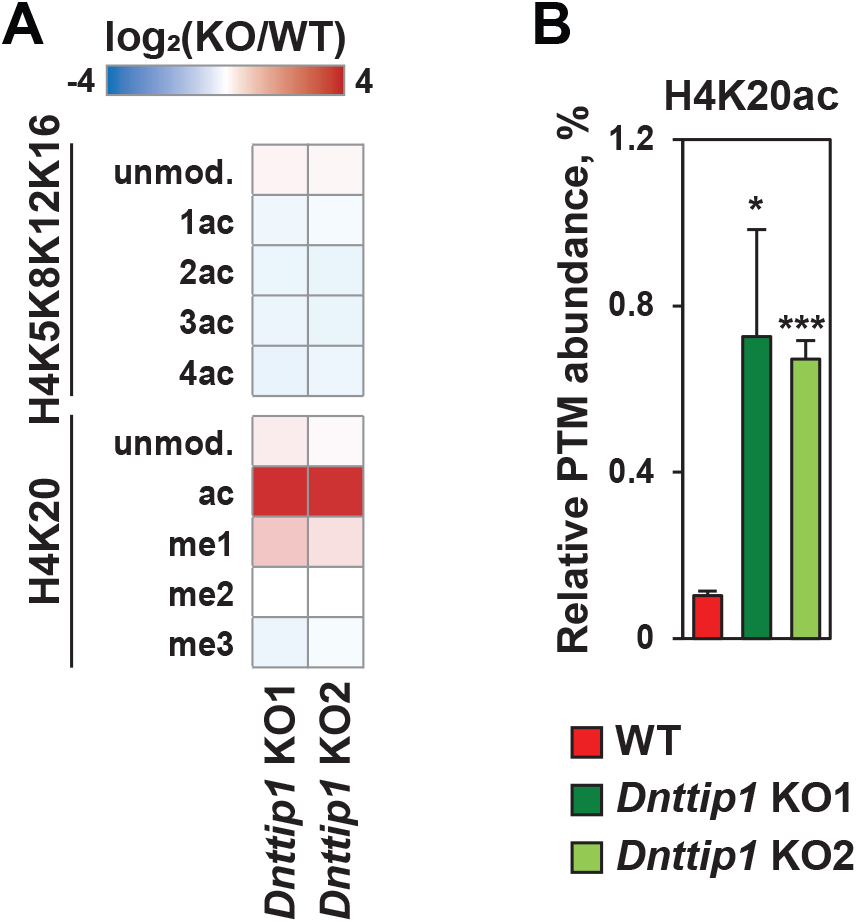
MiDAC is a global regulator of H4K20ac. **(A)** The heatmap shows the log_2_ fold change in the relative abundance of H4 post translational modifications (PTMs) between WT and two different *Dnttip1* KO mESC clones as assessed by mass spectrometry. **(B)** Mass spectrometry quantification of the relative PTM abundance of H4K20ac expressed as a percentage (%) in WT and two different *Dnttip1* KO mESC clones. Data are the means ± standard deviation of three independent experiments. Single and triple asterisks indicate adjusted p<0.05 and adjusted p<0.001, respectively (unpaired t-test, p values were corrected for multiple testing using the Benjamini-Hochberg method). H4K20ac levels are strongly increased in the absence of MiDAC function.

**Figure S5.**
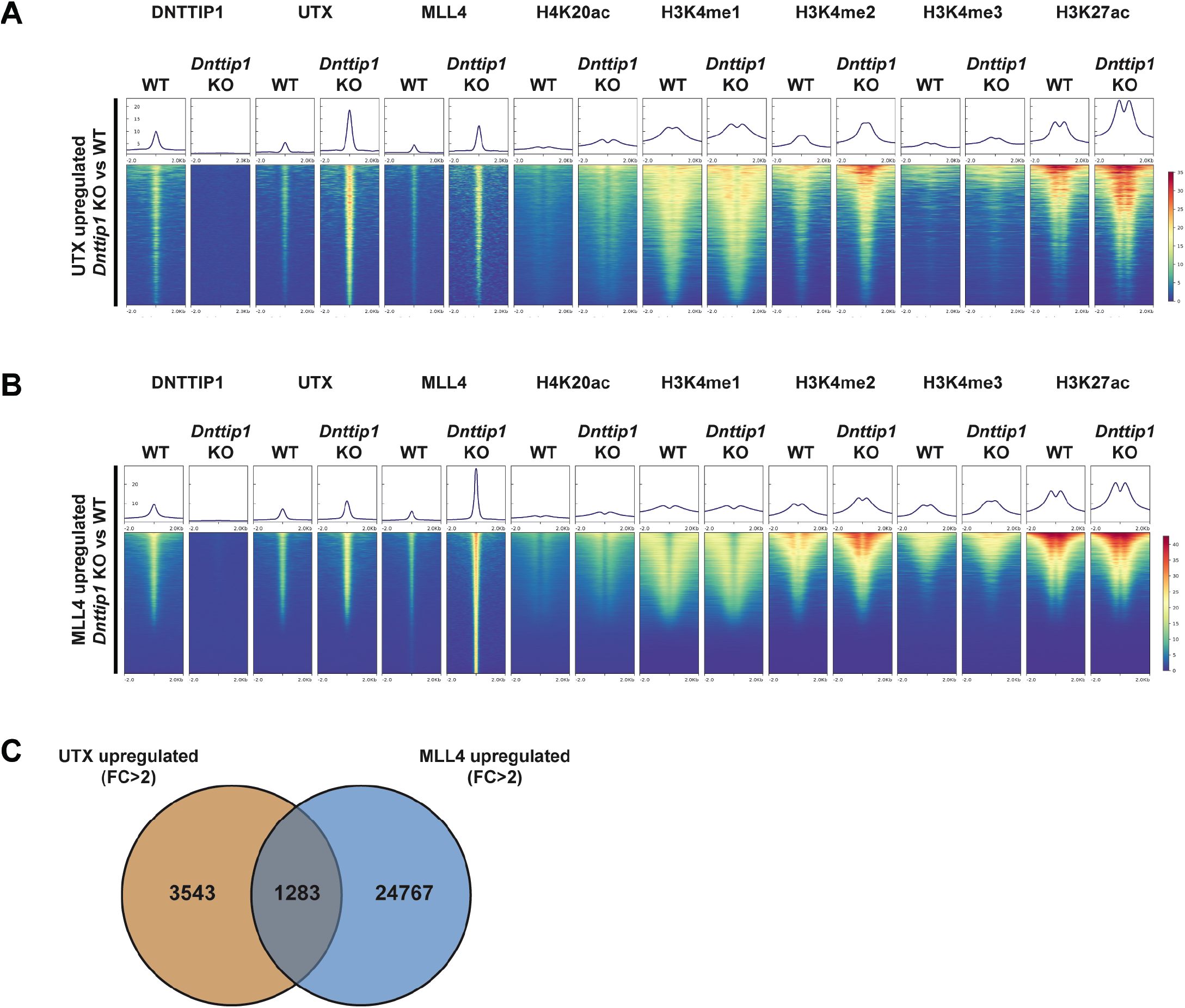
MiDAC negatively regulates the occupancy of UTX and MLL4 at regulatory elements genome-wide. **(A-B)** Heatmaps centered on 4,826 UTX upregulated peaks (FC>2) (A) and 26,084 MLL4 upregulated peaks (FC>2) (B) in *Dnttip1* KO compared to WT mESCs. Occupancy of DNTTIP1, UTX, MLL4, H4K20ac, H3K4me1, H3K4me2, H3K4me3, and H3K27ac in WT and *Dnttip1* KO mESCs is displayed. **(C)** Venn diagram depicting the overlap between UTX upregulated peaks (FC>2) and MLL4 upregulated peaks (FC>2) in *Dnttip1* KO versus WT mESCs.

**Figure S6.**
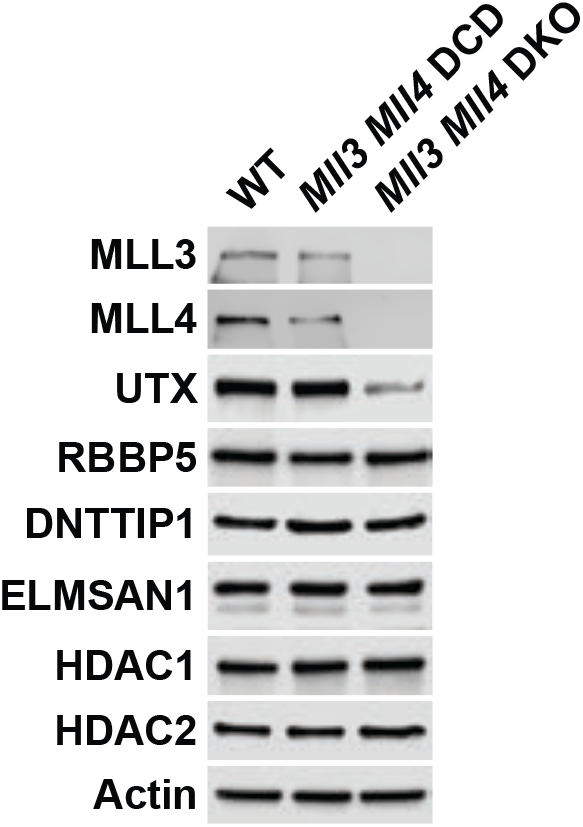
MiDAC stability does not depend on the MLL3/4 complexes. WB for various subunits of the MLL3/4 complexes and MiDAC from nuclear extracts of WT, *Mll3/4* DCD and *Mll3/4* DKO mESCs. As a result of MLL3/4 DKO, UTX protein levels are strongly decreased while protein levels of RBBP5 and MiDAC subunits (DNTTIP1, ELMSAN1, HDAC1, HDAC2) are unaffected in the absence of MLL3/4. Actin was used as loading control.

**Figure S7.**
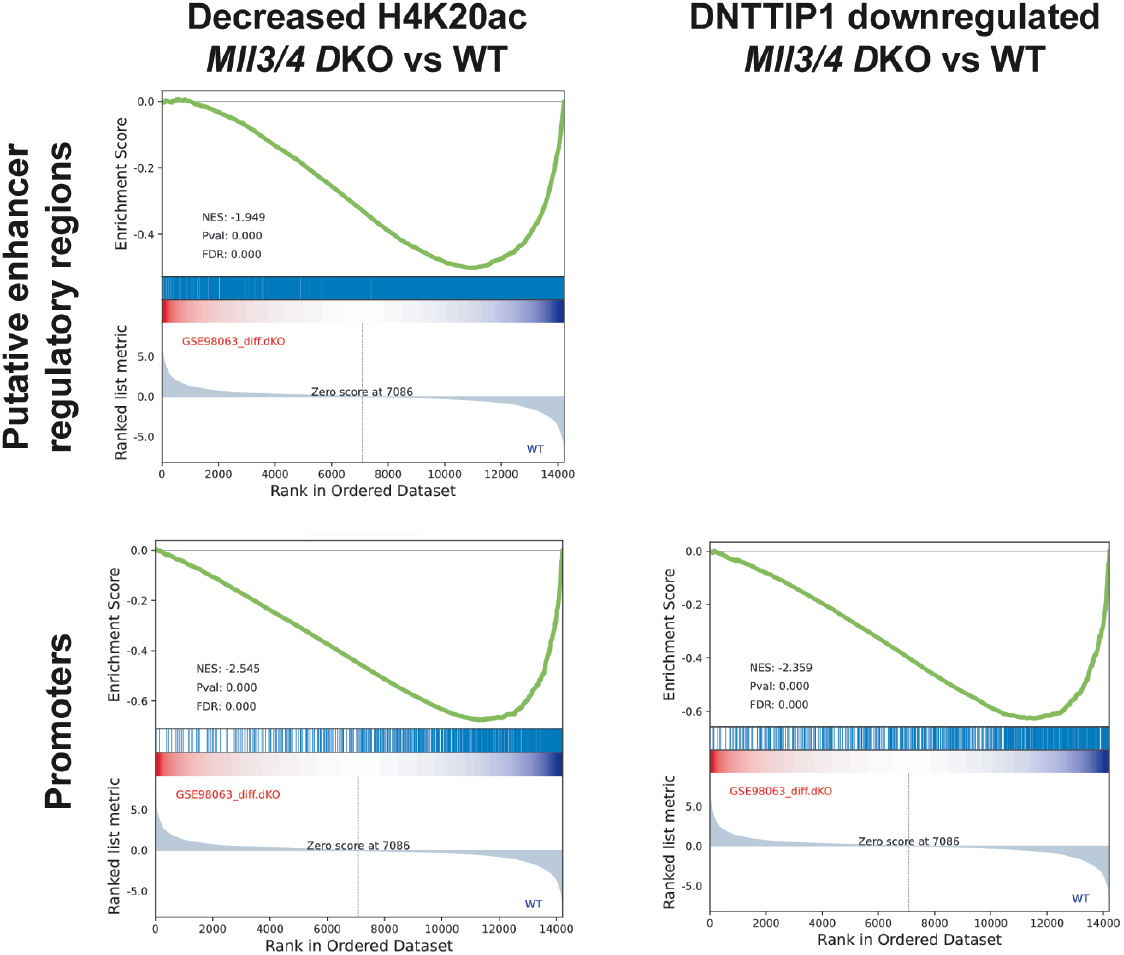
The loss of MLL3/4 results in a genome-wide decrease of H4K20ac and DNTTIP1 occupancy and is associated with gene repression. Gene set enrichment analysis (GSEA) showing transcriptional downregulation of genes associated with regions of decreased H4K20ac (FC<2) (left) and DNTTIP1 downregulated peaks (FC<2) (right) in *Mll3/4* DKO mESCs.

**Figure S8.**
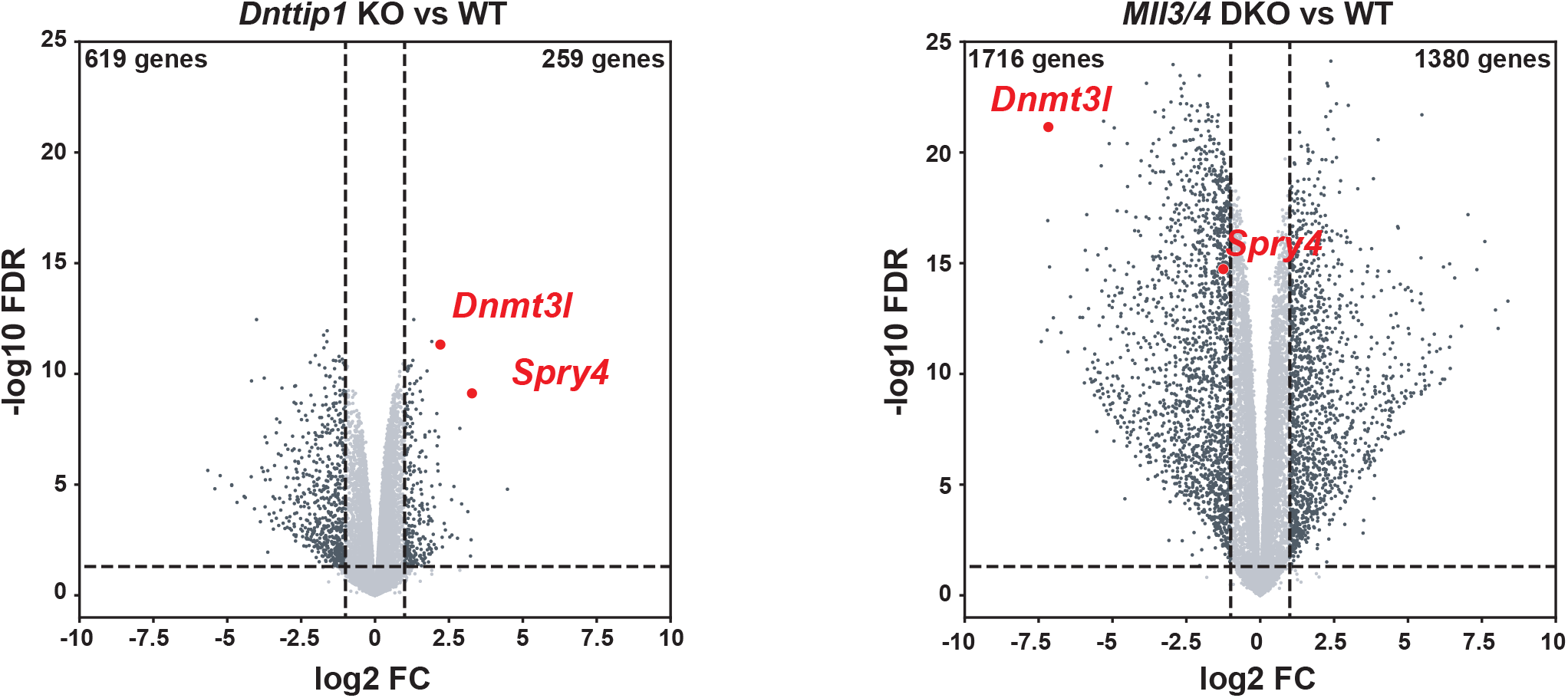
A specific gene expression program is controlled by the opposing functions of MiDAC and the MLL3/4 complexes. Volcano plots showing differentially expressed genes (FC>2 or <-2, p<0.05) in *Dnttip1* KO versus WT mESCs (left) and *Mll3/4* DKO versus WT mESCs (right).

**Table S2.**
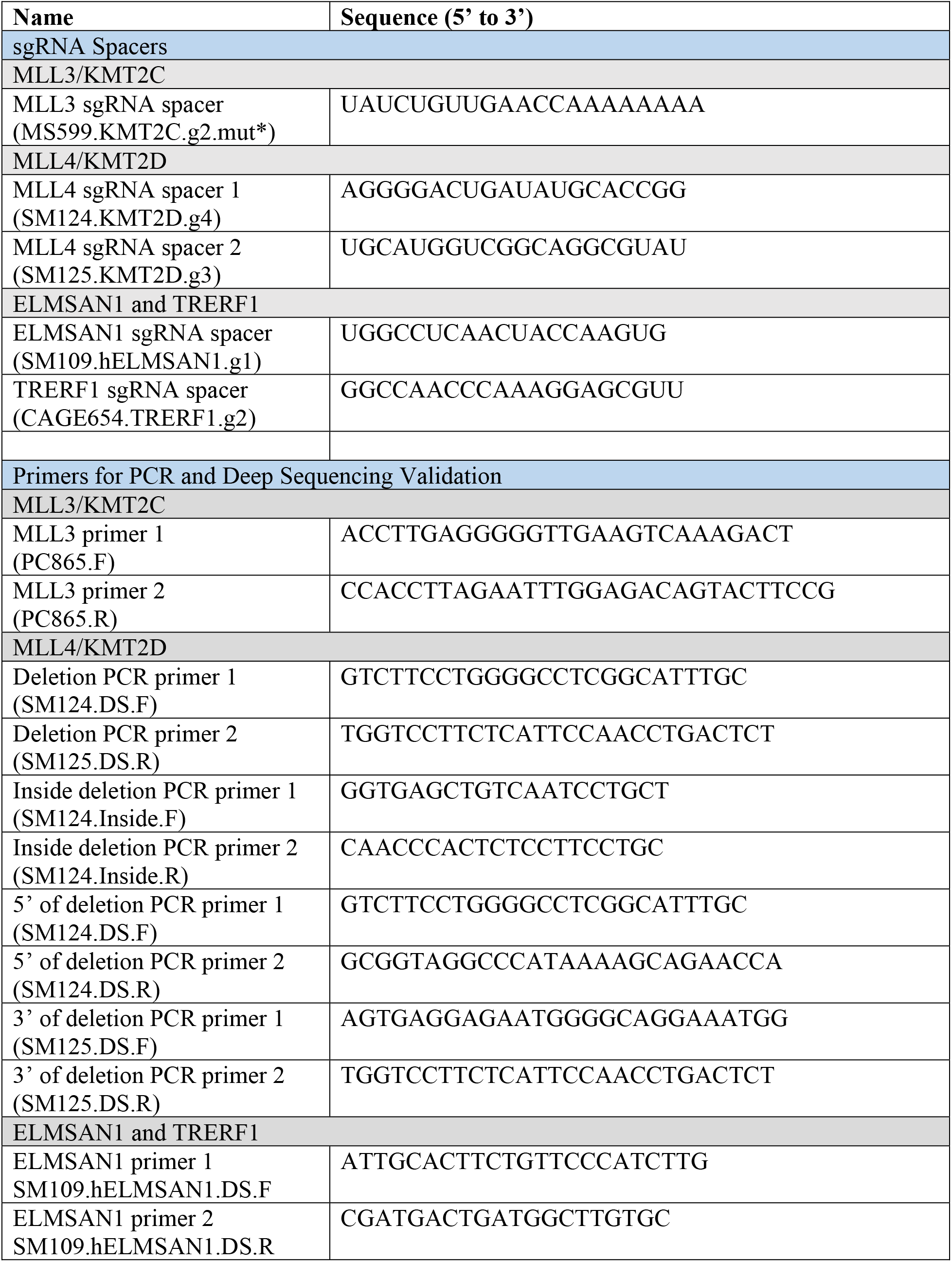
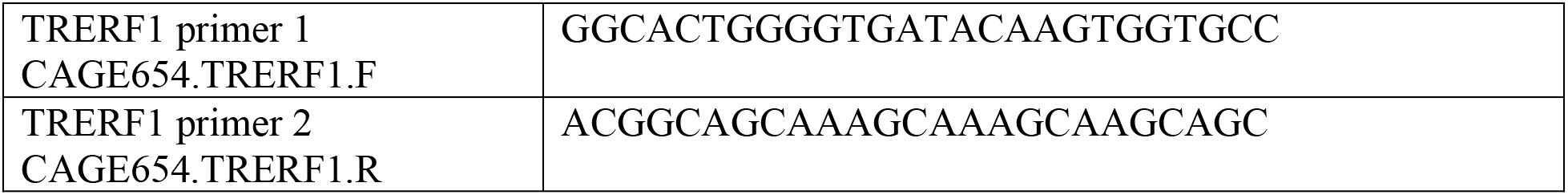
CRISRPR/Cas9 primers and editing construct sequences.

